# Activity dynamics allow early discrimination of infection-related survival outcomes

**DOI:** 10.1101/2025.04.21.649753

**Authors:** Megan A. M. Kutzer, Shima Abdullateef, Alejandro V. Cano, Iris L. Soare-Nguyen, Katy Monteith, Javier Escudero, Vasilis Dakos, Pedro F. Vale

## Abstract

Predicting transitions between health, disease, and death across biological systems remains an important challenge with significant implications for both ecological management and medical intervention. Although the principles underlying these transitions are increasingly recognised, accurate and tractable dynamical indicators of health-to-disease transitions remain rare, especially at the level of individual organisms. Here, we use dynamical statistical indicators of high-resolution activity time series to predict infection-related mortality. By analysing locomotor activity data from infected *Drosophila melanogaster* flies, we find that individual dynamical indicators, such as the mean, variance, autocorrelation, and permutation entropy, differed between flies that survived and those that died during the experiment. When these indicators were used to train a Random Forest model, the classifier performed well (AUC = 0.94), demonstrating an accuracy of 87.9% in discriminating between infected flies that would die from infection and those that would survive, with the strongest discriminatory power detected over 12 hours prior to death. Our findings show that combining these easy-to-compute, dynamical statistical indicators with machine learning enhances the ability to predict health deterioration in the *Drosophila* model. Conceptually, our findings emphasize that the integration of dynamical statistical metrics from physiological or behavioural time-series with machine learning approaches may offer a promising avenue for real-time health monitoring in both ecological and clinical settings.

**Highlights:**

- High-resolution locomotor activity time series distinguish infected flies that live or die
- Simple dynamical indicators (mean, SD, CV, lag-1 autocorrelation) jointly improve outcome discrimination
- Permutation entropy declines over time and is reduced in flies approaching acute infection-related death
- Random Forest models classify infection survival outcomes with high accuracy (AUC 0.94)
- Divergence in activity dynamics is detectable over 12-16 hours before death in flies that succumb to infection

## Background

Predicting abrupt transitions in a system’s state is a major challenge in ecological, earth and living systems [1]. It is difficult to anticipate abrupt changes in drylands shifting to a desert state [2], to detect the potential collapse of icesheets in West Antarctica [3], or to identify the episodes of depression in patients with psychological disorders [4]. In all these dynamical systems, there is usually a gradual insignificant change in the state of the system when even minute disturbances or slight changes in conditions may trigger abrupt shifts that lead to irreversible changes. These abrupt shifts, termed **“**critical transitions” or “catastrophic shifts”, are notoriously difficult to detect [1,5].

At a smaller scale, our ability to predict when individuals will transition between a state of health, disease, and death is poor for most biological systems [6–9]. Even in one of the best studied model systems of infection, the fruit fly *Drosophila melanogaster* [10], we struggle to predict when genetically identical flies given identical infections will either succumb to infection or develop asymptomatic infections [11–13]. Inaccurately predicting the death of a fly may seem trivial but failing to predict the progression of disease in a clinical setting may have more severe consequences. In general, predicting health deterioration is challenging even in cases where individuals are sick enough to be hospitalised [8,14].

A promising way that could help detect system transitions is based on the general observation that many dynamical systems tend to slow down prior to abrupt shifts [15]. This “slowing down” effect can be informative about future abrupt shifts in the state of any system [16], and it can serve as an early-warning signal for critical transitions [1]. Specifically, slowing down can be quantified through changes in the statistical metrics of a system’s temporal dynamics, such temporal changes in the mean, variance and autocorrelation of system states over time [17]. These slowing down indicators have been used in climate science, ecology, engineering, finance, and human physiology [18], and may have strong potential to be applied to chronic clinical conditions [6,8,9]. In addition to these slowing-down indicators, a distinct but commonly used metric to investigate a change in the state of dynamical systems is Permutation Entropy (PE). Unlike slowing down indicators that are meant to detect abrupt shifts, PE is a metric that is used to quantify the irregularity or disorder in the dynamics of a system [19,20]. The use of PE is based on the hypothesis that complexity, indicated by high entropy, is a hallmark of healthy physiological systems, and that a loss of complexity with illness and increased risk of death is associated with reduced entropy [7,20]. Numerous studies have demonstrated changes in entropy in various biological recordings, such as heart rate and brainwaves, in individuals with certain disorders [21–23].

In this study we evaluate whether generic critical slowing down (CSD) indicators and PE can serve as features for prognostic discrimination within infected individuals as they transition between health, disease, and death. We leveraged a fruit fly-bacterial infection model system where the underlying physiology and genetics of infection, immunity and behaviour are known in detail [10]. As a biomarker of health during infection, we focused on the locomotor activity of individual flies, measured continuously, at extremely high temporal resolution until death. Previous work has established that flies move in regular and repeatable crepuscular activity cycles when healthy, but show evidence of decreasing activity during infection as they approach death, especially in flies experiencing systemic or chronic infection [24–28]. Fly activity is therefore a useful biological signal to investigate transitions between life and infection-induced death, as it is a good proxy for host health and can be measured over time in the same individuals.

We further employed machine-learning (ML) approaches to test if we could discriminate between groups of flies that lived or died using the dynamical statistical metrics we calculated from activity time series. ML methods are data-driven decision tools used for pattern recognition, classification, or prediction tasks. The logic behind ML is that given a set of data, i.e. a training set, the algorithm aims to use patterns from the data itself and/or other user-defined inputs to perform a task, e.g. classify, predict, on a new set of data. ML approaches have started to be tested as tools for detecting transitions in both biological and mechanical systems [29,30]. Our results suggest that extracting dynamical statistical metrics from individual behavioural time-series, in combination with data-driven machine learning approaches, may offer a useful way to identify transitions between health and illness in organisms experiencing infection.

## Materials and Methods

### Collection of high-resolution activity data in infected flies

Our aim was to test whether we could identify differences in the behavioural time series of flies that lived or died following an infection. We collected high resolution activity data on female *Drosophila melanogaster* fruit flies, strain *w*^*1118*^ (Vienna *Drosophila* Resource Centre) using the Drosophila Activity Monitor [31].

To establish experimental infections, we used the generalist gram-negative bacterial pathogen *Pseudomonas aeruginosa* (strain PA14), a well-established model of infection in *Drosophila*. We inoculated 17 × 10 mL aliquots of sterile LB with a sterile inoculation loop from *P. aeruginosa* (PA14) stocks stored at -70ºC and incubated them at 37ºC 140 rpm overnight. The following morning, we reinoculated 17 × 9 mL fresh LB with 1 mL of each overnight culture and incubated them at 37ºC, 140 rpm until they reached exponential growth (between OD 0.6 – 0.8) approximately 1-2 hours later. The cultures were then spun down at 2500 × g at 4 C for 15 minutes until the supernatant was clear. After removing the supernatant, we resuspended the pellets in an appropriate volume of 1X Phosphate Buffered Saline (PBS) to reach an OD of 0.03. We used this bacterial culture to induce systemic infections in flies of line *w*^*1118*^ using a septic pin prick.

All experimental flies were females, raised at constant density for one generation, maintained at 25±1 °C on a 12 h: 12 h light: dark cycle at 60% humidity, on a standard cornmeal-yeast-sugar diet (14% protein; 1:6 protein: carbohydrate). Ten females and five males were placed in a vial for two days at 25°C, after which, the adults were discarded, and the offspring were left to eclose. Mated female offspring were collected at two to three days post-eclosion. We anesthetised females in groups of 8 using light CO_2_ and pricked them in the right lateral side of the thorax using a 5mm needle. This corresponded to a dose of approximately 200 cells per fly. The flies were placed back into the original vials to recover and then placed in Drosophila Activity Monitors (DAM5M) as described previously [32]. Prior to starting the experiment, we prepared DAM tubes by filling each with 2 cm of a 5% sucrose–agar solution. When loading the DAM monitors, flies were anaesthetized using CO_2_ and then placed individually into the tubes of each 32-tube activity monitor at random. The DAM monitors were kept in an incubator (25 °C on a 12 hr: 12 hr light:dark cycle, 60% relative humidity) to minimize possible disturbance and disruption, and each monitor contained either one empty tube or no tube as negative controls. Each monitor was connected to a laptop, running the DAMSystem3 data collection software, which was placed in the incubator. Activity counts and lifespan were recorded in 1-minute bins for two weeks, after which we checked individual survival in each tube by eye.

### Data pre-processing

Data from the DAM were processed in R. The raw data obtained from the DAM monitors contained a single file for each monitor, which included an activity time series for a total of 32 flies per monitor. After concatenating the data files for each monitor, we took the last recorded movement as the time of death. We differentiated between inactivity and death in the instances where the raw data recorded “no movement,” originally represented as zeros. Non-movement due solely to inactivity, such as periods of rest or sleep, were retained as zeros. Conversely, we scored a fly as dead if there were no recorded movements in 1000 minutes and regular movements never resumed. These occurrences were then marked as “Not a Number” (NaN) within the dataset to distinguish them clearly from periods of inactivity. The last real movement was recorded as the time of death. Following this pre-processing step, we retained flies from the analyses (250 determined as “dead” at some point during the 14-day experimental period, and 161 that while also infected, remained “alive”. We use these terms to distinguish between these two groups throughout the rest of the text). The experiment contained 26 sham-pricked controls. These were removed from our subsequent analyses and are shown in Figure S1.

### Estimation of dynamical statistical indicators of fly activity

We evaluated 3 statistical metrics along fly activity time series typically used as dynamical indicators of abrupt transitions: namely trends in variance (as standard deviation, SD), coefficient of variation (CV, defined as the ratio of the SD to the mean, where a higher CV, indicates greater dispersion around the mean) and autocorrelation at-lag-1 (AR1, linear relationship between consecutive lagged values in a time series (values between -1 and 1) where a value of 1 indicates a perfect positive correlation). We estimated these three indicators within rolling, non-overlapping 60-minute windows from the beginning to the end of each time series. We then computed the change in the indicators using a Kendall’s *τ* rank correlation of the estimated values from each window. Kendall’s *τ* is a non-parametric statistical measure that determines the strength and direction of association between variables. Kendall’s *τ* takes values between -1 and 1, implying a decrease or an increase in SD, CV or AR1 respectively. We also estimated trends in the mean fly activity using the same rolling windows. In addition to trends, we computed SD, CV, AR1 and mean fly activity at the scale of the total time series. Metrics computed at the scale of the total time series were rescaled to 0-1 by subtracting the minimum value from each observation and dividing by the range; this was done only for visualisation purposes. We statistically compared these 8 metrics between “dead” and “alive” fly groups using linear mixed models using either the lme4 package [33] or the nlme package [34,35], depending on variance imbalance. P-values were corrected for multiple testing using the Benjamini and Hochberg method. (Appendix 1).

### Permutation entropy

Permutation entropy quantifies the predictability of a time series by assessing the relative ordering of values within successive windows. Given a time-series of length *N*, where data is represented as *x* = {*x*_1_, *x*_2_, … *x*_*n*_} and specified parameters for the embedding dimension *m* and time delay *τ*, vectors of length *m* with a delay of *τ* are constructed as *X*_*i*_ = {*x*_*i*_, *x*_*i* + *τ*_, … *x*_*i*+(m−1) *τ*_}. The permutation of each vector *X*_*i*_ is determined by arranging its elements in ascending order. Permutation entropy is then calculated using the formula:

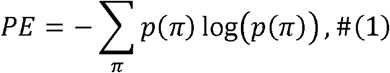

where *p*(π) represents the probability of occurrence of each distinct permutation π. For a detailed mathematical theorem of permutation entropy, readers are referred to [20,36]. In this analysis, a 30-minute non-overlapping window was used. The PermEn function from the EntropyHub toolbox [37] in MATLAB was used to calculate permutation entropy. The embedding dimension *m* was set to 3 to achieve robust estimations of the probability of occurrence of each permutation π. The time delay *τ* was set to 3 and 1 to maintain the high temporal resolution of the data. These parameters are in line with common practice in the literature [38]. All entropy computations were performed using MATLAB R2022b, a programming language and numeric computing environment commonly used in nonlinear analysis and signal processing. Differences between the Permutation Entropy in each group were assessed using linear mixed models detailed in Appendix 1.

### Random forest model for classifying dead and alive flies

Pre-processed time series data of fly activity were analysed to predict mortality. A Random Forest model was trained to classify flies as either “alive” or “dead” based on the calculated metrics. We used mean activity, standard deviation (SD), coefficient of variation (CV), and lag-1 autocorrelation (AR1) as predictors, computed either across the full time series or within aligned pre-death windows. Permutation entropy was analysed separately; including PE as an additional predictor did not materially change performance, so we report the more parsimonious model. The dataset was split into training (60%) and testing (40%) sets using stratified random sampling, ensuring a representative distribution of “alive” and “dead” flies. We then examined the Out Of Bag (OOB) error rate to determine the optimal number of trees for our random forest models using the randomForest package in R. We tested our models using 300 trees. After training the models, we tested accuracy using the test datasets and examined the confusion matrices and associated statistics, i.e. Kappa (how much better the classifier performs than would be expected by chance), McNemar’s test p-value (tests for balance between false negatives and false positives: if p is not significant there is no evidence of an imbalance), sensitivity, specificity, balanced accuracy, using the confusionMatrix function in the “caret” package [39]. We checked variable importance using the importance and varImpPlot functions in the “randomForest” package [40]. To further evaluate the model’s discriminatory power, a Receiver Operating Characteristic (ROC) analysis was performed by plotting the true positive rate against the false positive rate. The area under the ROC curve (AUC) was calculated using the “roc” function in the pROC package [41] as a measure of the model’s ability to distinguish between “alive” and “dead” flies. A ROC curve shows how well a model can distinguish between two groups. It plots the trade-off between correctly identifying true positives and falsely classifying negatives as positives, so the closer the curve is to the top left corner and the larger the AUC, the better the model is at discriminating between the groups. An AUC of 0.5 means that model is no better at distinguishing groups than random. Random forest models and validation steps are available in Appendix 1 as an R Markdown. All raw data and analysis code is publicly available at https://doi.org/10.5281/zenodo.20055750.

## Results

### High resolution activity time series differs according to infected survival outcome

Time series data were collected on 411 infected flies. Of those infected flies, 161 (∼40%) survived until the end of the experiment, while 250 (∼60%) died (Figure 1A). We also collected data on 26 sham-pricked uninfected flies; among uninfected flies, 22 survived and 4 died (Figure S1). Hourly mean movement data revealed distinct activity patterns between infected flies that survived and those that died during the experiment (Figure 1B). Flies that remained alive exhibited consistent rhythmic activity throughout the time series, characterized by pronounced peaks and troughs in movement due to consistent circadian cycles (Figure 1B, top panel, blue). In contrast, flies that died showed a progressive decline in activity over time, with reduced rhythmicity and lower overall movement levels (Figure 1B, bottom panel, orange). Infected flies that survived the 14-day experimental period maintained higher and more stable mean activity compared those that died (Figure 1B). These results suggest a potential link between movement patterns and survival outcomes in infected flies.

**Figure 1.**
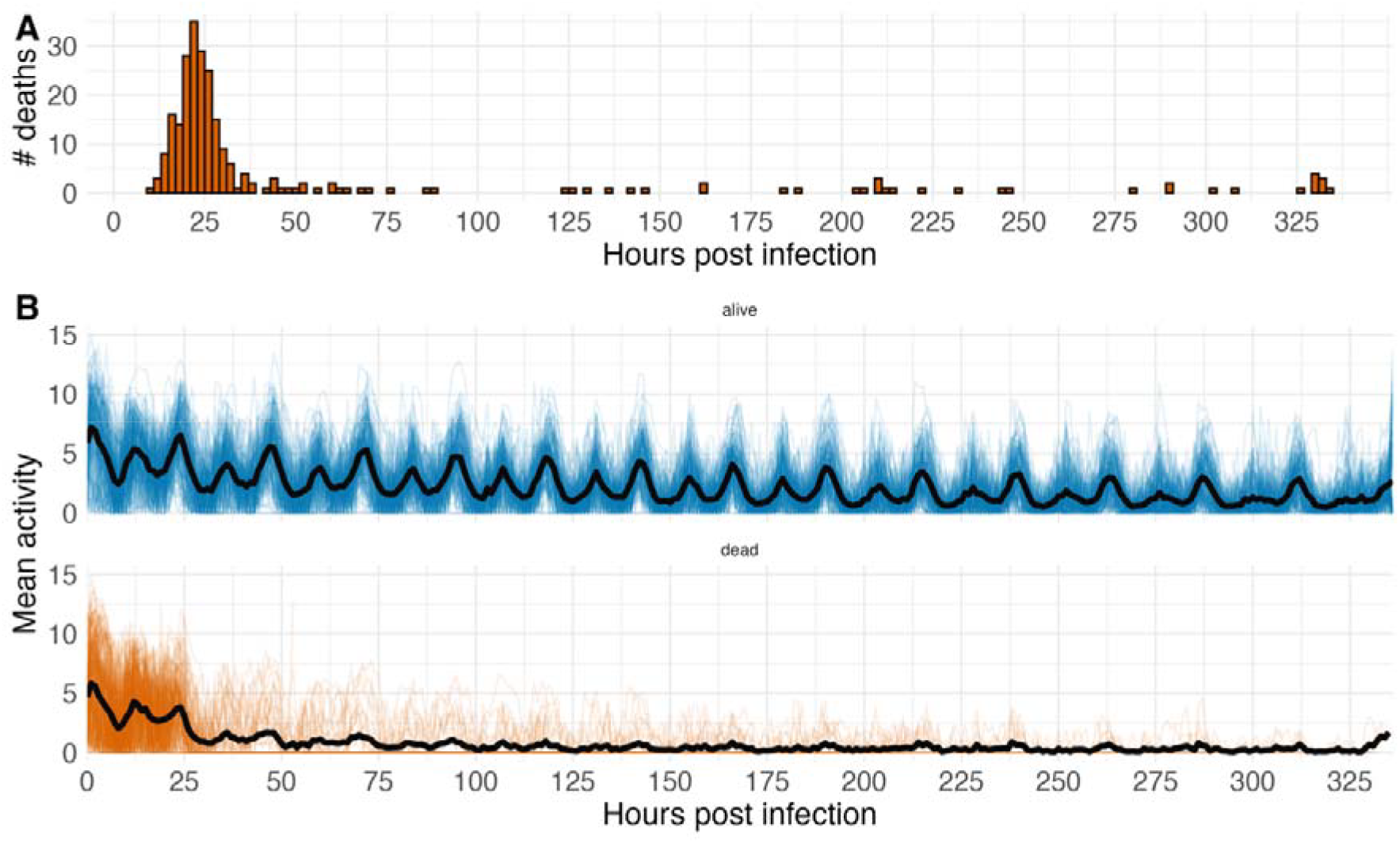
Activity differs in infected survival outcomes. A. Number of deaths per hour (non-cumulative) over 14 days post infection. Deaths from systemic infections at this dose tend to occur within 36 hours. B. Time series showing the changes in hourly mean movement over time for infected flies that lived during the experiment (top panel, blue), and flies that died (bottom panel, orange). The black line indicates averages in each panel.

### Individual statistical metrics are modest discriminators of flies that live or die

When calculating individual dynamical metrics on each time series we found a reasonable ability to distinguish between flies that survived and those that died (Figure 2). Metrics calculated using the entire time series (Figure 2A) were aggregated by status (alive or dead) and displayed as percentile ranks. We observed significant differences between alive (blue) and dead (orange) flies for all metrics (Figure 2A). Trends in the metrics calculated using a nonoverlapping rolling window of 1 hour (Figure 2B) and represented as Kendall’s correlation coefficients revealed significant differences only for CV and SD. Given the high variance observed in all metrics for both classes, these findings suggest that individual dynamical metrics alone may not reliably discriminate between survival outcomes in infected flies.

**Figure 2.**
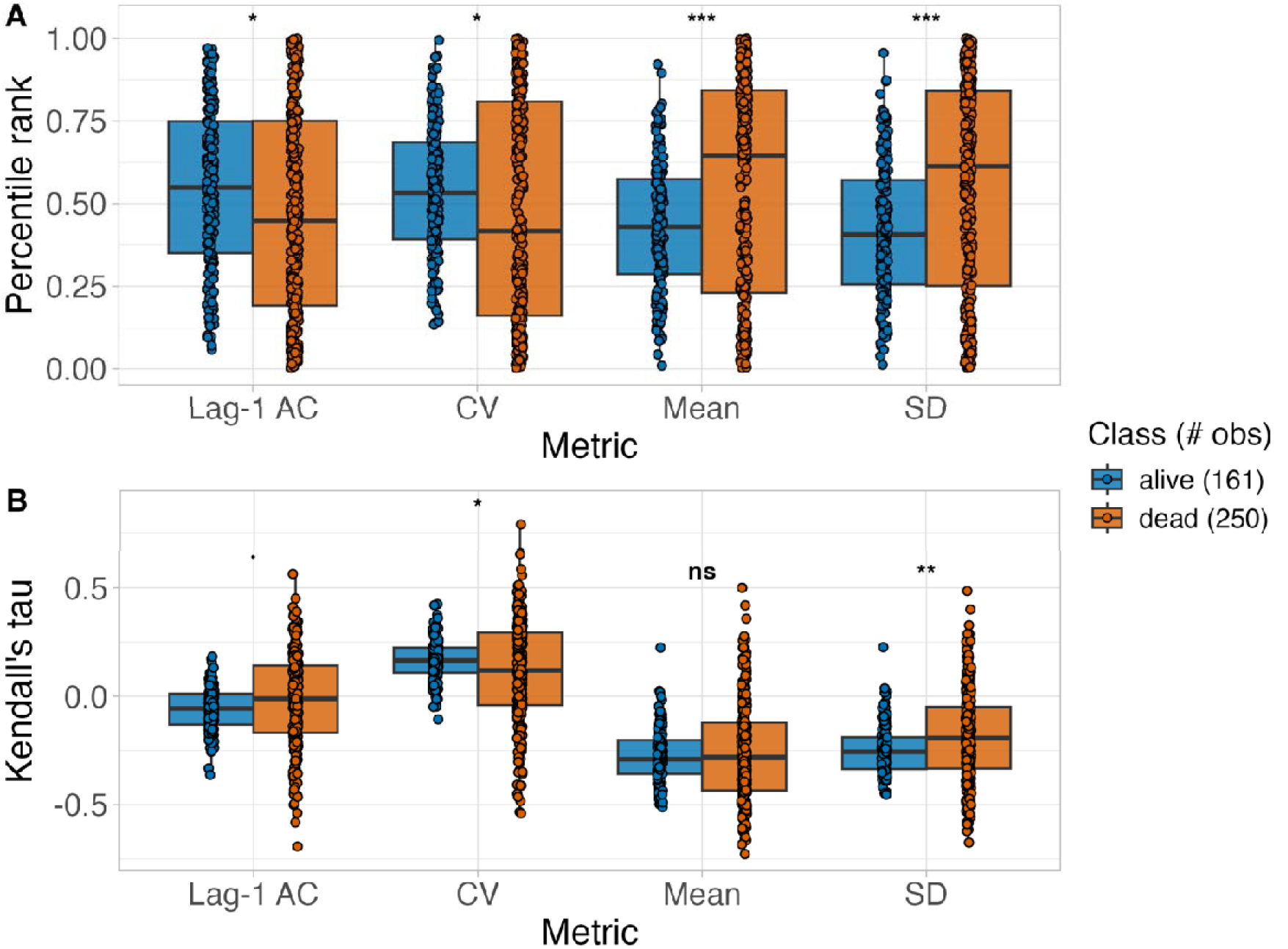
Individual dynamical metrics discriminate between dead and alive classes. Boxplots show the values of all individual metrics, aggregated by class. A) Metrics calculated using the entire time series, and for visualisation purposes, the Y-axis shows the percentile rank of each metric. B) Trends in the metrics calculated using consecutive 1 hour activity windows The Y-axis shows Kendall’s where a positive value indicates an increase in the metric and a negative value a decrease. Statistically significant differences are indicated with stars (Benjamini-Hochberg corrected p-values, LMMs).

### Permutation entropy declines over time regardless of survival class

In addition to the generic statistical indicators that are commonly used to detect critical transitions in dynamical systems, we also calculated the permutation entropy of each time-series. Permutation entropy provides a measure of the unpredictability of a time series, which can be inversely correlated with system health [20]. As system behaviour becomes more predictable and less complex, this often signals a deterioration or loss of function [7]. We calculated the permutation entropy from the start of each fly time-series until the last movement of each fly (or for 14 days in flies that remained alive) in 30-minute non-overlapping windows (Figure 3). In both sets of flies that survived and died from infection, the average permutation entropy initially shows regular oscillatory patterns that reflects the underlying circadian rhythm of fly activity, although PE in the ‘dead’ class appears to be declining in the period preceding death (Figure 3A). These regular cycles show higher entropy during periods of fly activity, and lower entropy during periods of inactivity, when activity pattens become more predictable. There is also a general decline in permutation entropy between the start and end of the experiment (Figure 3C). This suggests physiological function declines in both sets of flies, which is expected both as a cause of infection, and because the conditions inside a narrow DAM tube are likely to cause some deterioration in the health of each fly.

**Figure 3.**
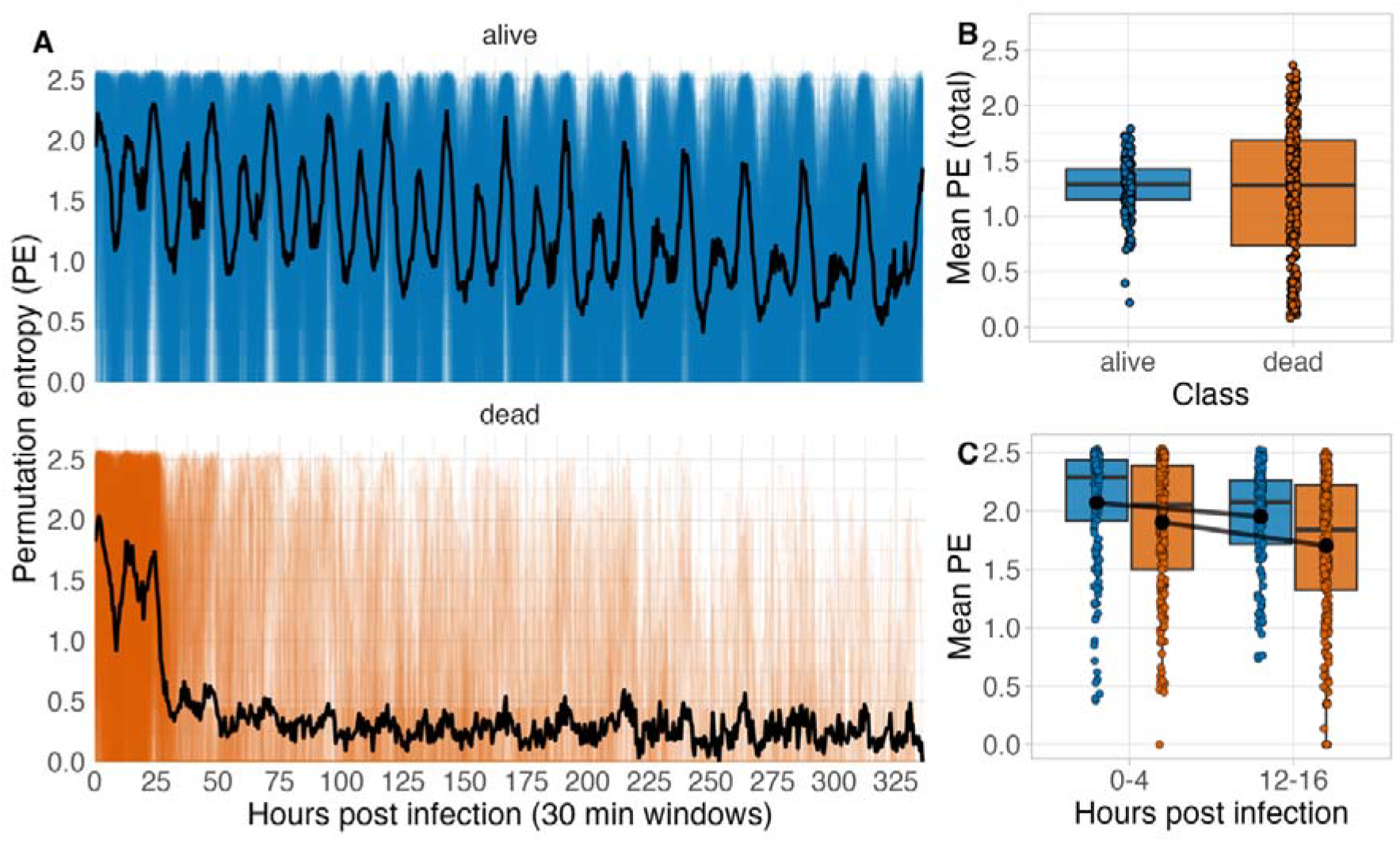
Permutation entropy declines with age and is lower in individuals approaching death by acute infection. A – Top panel. The aggregated PE time series for the surviving flies from the infection experiment compared with all PE time series exhibiting the same characteristics, Bottom panel - The aggregated PE time series for the deceased flies from the infection experiment (aligned to the time of death). B. The mean and distribution of PE during the whole recording. There is no difference in PE between classes (mean PE dead = 1.21, mean PE alive = 1.27; LMM, Class: p = 0.229). C) The mean and distribution of PE in the first four hours post infection versus 12 to 16 hours post infection. Flies that survive the experiment (alive, blue) exhibit significantly higher permutation entropy on average than flies that die (orange) from infection during the acute infection period (LMM, Class: p<0.00001).

Averaged across the length of all individual time series, there was no difference in permutation entropy in flies that died compared to those that survived, averaged over the entire time series (Figure 3C; mean PE dead = 1.21, mean PE alive = 1.27; LMM, class: p = 0.229). However, we noticed that PE appears to decline as flies approach death, so we compared the first four hours post-infection to 12-16 hours post-infection, as the majority of the “dead” individuals were alive in the latter window. We found that dying flies tend to have lower PE overall than flies that survive the experimental period (LMM, class: p<0.0001), consistent with the hypothesis that higher entropy reflects the greater complexity of healthier systems, whereas individuals approaching death show a loss of complexity.

### Random Forest models uncover dynamical differences between flies that live and die several hours before death

While individual dynamic metrics alone did not strongly discriminate between infected flies that lived or died, we tested if we could improve this discrimination using a data-driven, machine-learning approach by including information from multiple metrics in a Random Forest (RF) model.

We ran two types of RF model. In one approach, the Mean, SD, CV and autocorrelation were calculated for the entire length of each fly activity time series and used as predictors in the RF model. This model showed strong predictive performance (AUC = 0.943, Figure 4A), and classified survival outcome with 87.88% accuracy (95 % CI: 81.9-92.4%). The confusion matrix indicated that the model correctly identified 92.9% of flies that died (sensitivity) and 80.3% of flies that survived (specificity), with a balanced accuracy of 86.6% (Figure 4A). Agreement between the predicted and observed outcomes was also substantial (k = 0.744). McNemar’s test was not significant (p = 0.264), indicating that the model did not show strong asymmetry in the two types of classification error.

**Figure 4.**
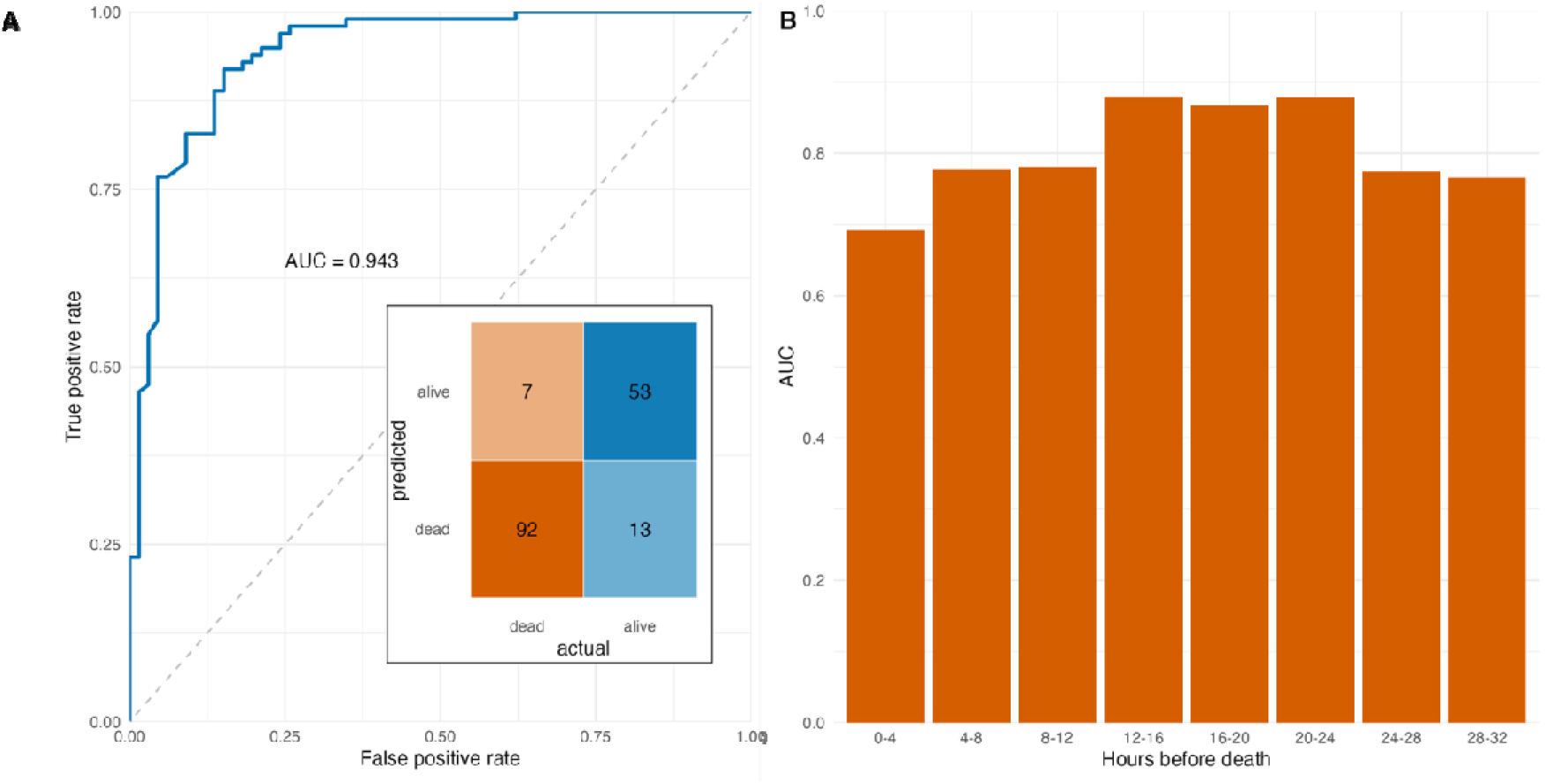
Random Forest models predict survival outcome at least 12h prior to death. A) The area under the ROC curve (AUC) represents the probability that the model, if given a randomly chosen positive and negative example, will rank the positive higher than the negative. The inset confusion matrix summarises the counts of true positives, true negatives, false positives, and false negatives predicted by the RF. Both the AUC and the confusion matrix refer to the model on the full time series. B. AUC for each 4-hour time window (hours preceding death) from a model trained on successive 4-hour windows preceding the death of each fly.

Next, we ran a second RF model to ask how early differences in activity dynamics could be detected before death. Because each fly died at a different time, in this RF model we aligned all time-series by the last recorded movement so that activity dynamics could be compared across equivalent time intervals preceding death. We estimated mean, SD, CV and AR1 in 4-hour windows preceding an individual’s last movement, up to 28-32 hours before death, as over 50% of flies had died before 32 hours post infection (1.3 days, Figure 1A). If a fly did not survive long enough to contribute data to a given window, it was excluded from the analysis for that window. We found better classification ability than by chance for all windows (Figure 4B), with the strongest performance in the 12-16 hour window before death (AUC = 0.88, Figure 4B). In this window, the model correctly classified 85% of the flies that died and 75% of the flies that survived, demonstrating an overall accuracy of 81.1% and indicating that mortality associated changes in behavioural dynamics were already detectable well before death.

## Discussion

We set out to test whether easy-to-compute, generic dynamical statistical indicators drawn from high-resolution locomotor activity could forecast infection-related mortality in individual flies. Our analysis revealed three major results. First, activity dynamics of infected flies that ultimately died differed from those that survived: when computed across the whole time series, mean activity, variance (SD), coefficient of variation (CV), and lag-1 autocorrelation (AR1) all differed significantly between classes, albeit with substantial overlap in their distributions. Second, when we aligned activity to time-of-death and asked how early differences became detectable, a Random Forest model built from these indicators achieved high discriminatory performance, with the strongest signal 12–16 hours before death. Third, permutation entropy (PE) declined over time across all flies and was lower in flies that died during the acute phase but did not differ between classes when averaged across the whole experiment. Together these results show that activity carries prognostic information, but that the dynamical signatures do not conform neatly to the canonical expectations under critical slowing down (CSD) or to a large, class-wide reduction in entropy.

The ability to predict dynamical transitions using dynamical statistical metrics in a variety of systems is appealing because it suggests that there are consistent changes in the dynamics of a system that can be informative about future abrupt shifts in the system state [1,16,18]. For example, this approach has been employed to estimate slowing down indicators for critical transitions in large scale systems such as climatic shifts [42] or to understand the behaviour of populations following a perturbation [43]. However, our results do not completely align with critical slowing down (CSD). Under the CSD framework, as a system approaches a critical transition it should exhibit slower recovery rates from perturbations, which are reflected statistically as rising variance and rising autocorrelation (AR1), sometimes accompanied by increasing CV [1,16,17]. If infection-related death in flies reflected such a transition, we would therefore expect upward trends in these indicators in the pre-death period. Trends estimated from rolling windows showed significant class differences for SD and CV but not for AR1, and the dominant pattern in flies that died was a decline in activity amplitude and rhythmicity rather than a rise in autocorrelation. When these indicators were calculated across the full time series, we observed that means differed significantly but with broad overlap, indicating that single indicators were not, on their own, strongly diagnostic. Therefore, the absence of increasing AR1 and variance suggests that this acute infection setting is likely not a classical CSD process.

It is unclear whether this finding is a general feature of infections, or specific to this host-pathogen system. While applying slowing down indicators to infection-induced health transition remain rare, in some cases they have been applied to identify warning signals in the context of infection. For example, time-series analysis of interleukin-6 (IL-6) concentration levels in pigs before and after infection with *Actinobacillus pleuropneumoniae* found that increases in the amplitude of IL-6 fluctuations were significant warnings of infection, suggesting that rising fluctuations rather than absolute levels of IL-6 were more predictive of the health-to-disease transition [44]. This result has led to speculation that variability in specific pro-inflammatory cytokines in human subjects may be used to monitor possible critical transitions in inflammatory responses [8,9], but to our knowledge this hypothesis has not been tested.

We also investigated if the state of a fly can be predicted by the PE, which summarises information about the predictability of a time series to identify more gradual transitions in the health state of several dynamical systems. Permutation Entropy has resulted in promising results in the analysis of sleep stages from electroencephalogram (EEG) [45] and electrocardiogram (ECG) data in the context of cardiovascular health [23], and neuroimaging MRI data in early Alzheimer’s disease [46]. Specifically, the loss-of-complexity hypothesis posits that healthy physiology exhibits richer, more irregular dynamics (higher entropy), with illness and ageing associated with simplification and reduced entropy [7,20]. Consistent with this, PE declined over the course of the experiment in both groups of flies (indicating a general decay in physiological performance while in the DAM tube), and in the acute post-infection window dying flies exhibited lower PE than survivors. However, when averaged over the entire recording, differences between flies that lived and those that died were not detectable.

Although individual dynamical indicators showed only modest discrimination between flies that lived or died during infection, jointly they yielded strong discrimination, both across the full time series (AUC =0.94) and, critically, within pre-death windows up to 12-16 h before death (AUC =0.88). We interpret this improvement under RF as the mean, SD, CV and AR1 capture partially independent aspects of activity dynamics over time: general reduction of movement (mean), loss of rhythmic amplitude (SD), disproportionate variance relative to the declining mean (CV), and consequently, lower predictability of movement between sequential timepoints (AR1). Their combination therefore likely provides a composite signature of health deterioration that is detectable earlier than any one metric alone. From an applied perspective, this is encouraging, as relatively simple, interpretable features extracted in near-real time could provide prognostic information without requiring full knowledge of the underlying transition type.

Applying ML, particularly random forest RF models, to the prediction of health conditions based on behavioural and physiological time series data has garnered extensive interest in both the medical, ecological, and biotechnological fields [29,30]. Random forests are particularly suited for these tasks due to their ability to handle large datasets with numerous variables, manage unbalanced data, and to provide insights into feature importance. For example, using high resolution physiological data, RF models have been employed to identify individuals with cardiovascular disease [47,48], disordered sleep patterns [49], or depression and anxiety [50,51]. Features from such time series can be included as predictors to forecast the likelihood of patient hospital readmission, which can aid in improving personalised care plans [52,53].

In summary, our results show that high-resolution behaviour contains a robust, early signature of infection-related mortality, with the strongest signal emerging more than half a circadian cycle before death. Permutation Entropy supports a general loss of complexity with time and a more pronounced decline in flies that die during the acute infection phase. This integrated approach provides a framework for applying these methods to the deterioration of health in individuals. We have based our analyses on the premise that behavioural time-series reflect changes in health to the degree required to identify future mortality outcomes. In the case of fruit flies, this is supported by work that has also identified clear shifts in activity patterns following bacterial or viral challenge [25,27,28,32,54,55]. Expanding this idea to other organisms would require that the phenotype being measured is an accurate biomarker of individual health, and that it can be measured repeatedly in the same individual at high resolution during the transition between health, disease, and death. The integration of dynamical statistical metrics from physiological time-series with machine learning approaches may offer a promising avenue for predicting transitions from health-to-disease in a variety of ecological and clinical environments.

## Acknowledgements

We thank Md Kabir Ahasanul and Tameem Somalya for their initial exploration of this data. This work was funded by a Royal Society International Exchange grant IES\R2\222092 awarded to PFV and VD, and a School of Biological Sciences Seed Fund (University of Edinburgh) awarded to PFV and JE. For the purpose of open access, the authors have applied a Creative Commons Attribution (CC BY) licence to any Author Accepted Manuscript version arising from this submission.

---

**Figure S1:**
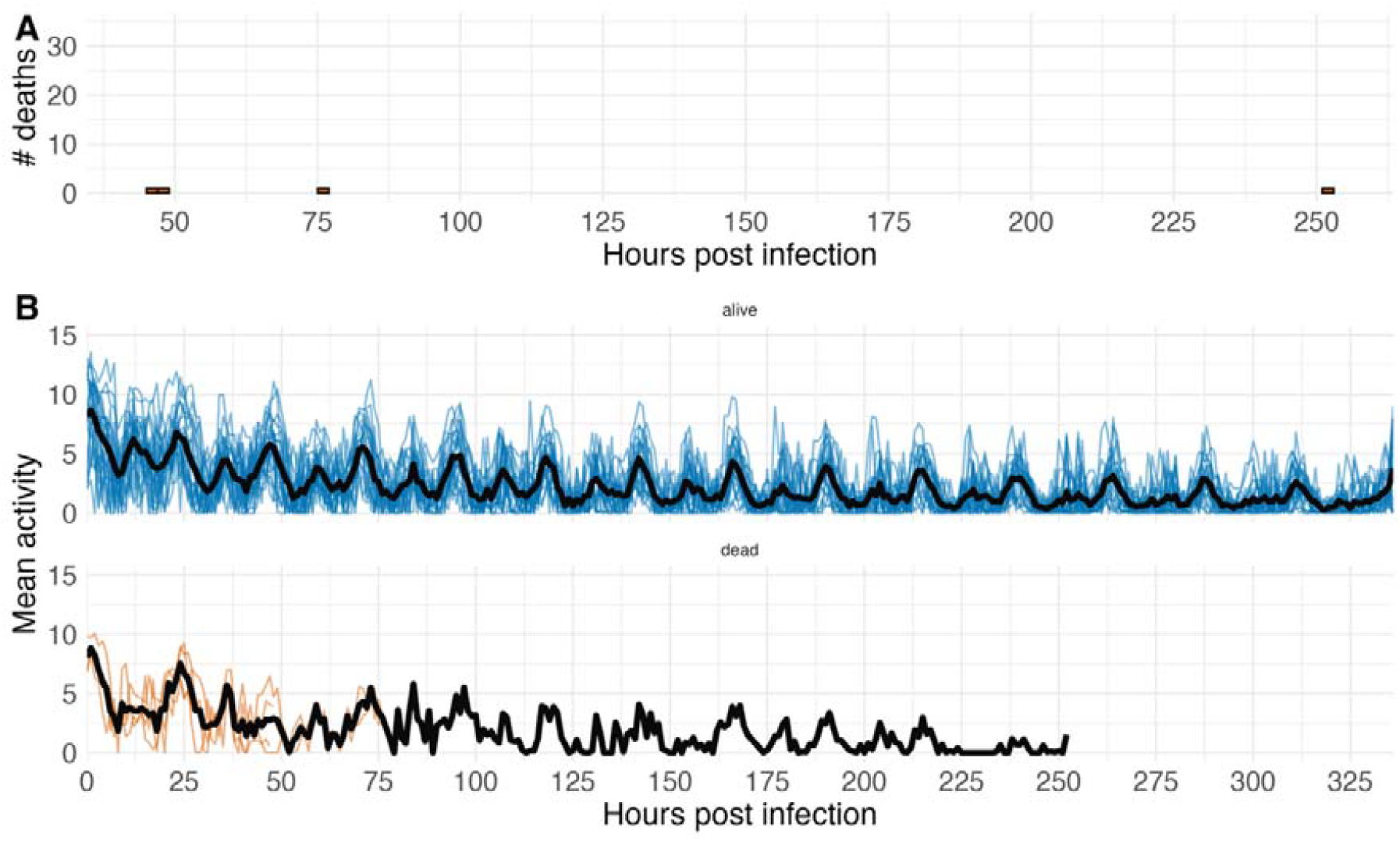
Uninfected controls. A) A total of four out of twenty-six uninfected flies died during the experimental period. B) Time series showing the changes in hourly mean movement over time for uninfected flies that lived during the experiment (top panel, blue), and flies that died (bottom panel, orange). The black line indicates average activity of alive flies in each panel.

