## Appendix for "Activity dynamics allow early discrimination of infection-related survival outcomes": Appendix-1.html

Critical transitions markdown


### Critical transitions markdown

#### 2026-04-24

### Load libraries and define functions

```
## 
## Attaching package: 'dplyr'
```

```
## The following objects are masked from 'package:stats':
## 
##     filter, lag
```

```
## The following objects are masked from 'package:base':
## 
##     intersect, setdiff, setequal, union
```

```
## 
## Attaching package: 'tidyr'
```

```
## The following object is masked from 'package:reshape2':
## 
##     smiths
```

```
## Warning: package 'ggplot2' was built under R version 4.5.2
```

```
## randomForest 4.7-1.2
```

```
## Type rfNews() to see new features/changes/bug fixes.
```

```
## 
## Attaching package: 'randomForest'
```

```
## The following object is masked from 'package:ggplot2':
## 
##     margin
```

```
## The following object is masked from 'package:dplyr':
## 
##     combine
```

```
## Type 'citation("pROC")' for a citation.
```

```
## 
## Attaching package: 'pROC'
```

```
## The following objects are masked from 'package:stats':
## 
##     cov, smooth, var
```

```
## Loading required package: lattice
```

```
## 
## Attaching package: 'rstatix'
```

```
## The following object is masked from 'package:stats':
## 
##     filter
```

```
## Loading required package: survival
```

```
## 
## Attaching package: 'survival'
```

```
## The following object is masked from 'package:caret':
## 
##     cluster
```

```
## 
## Attaching package: 'coin'
```

```
## The following objects are masked from 'package:rstatix':
## 
##     chisq_test, friedman_test, kruskal_test, sign_test, wilcox_test
```

```
## Warning: package 'lme4' was built under R version 4.5.2
```

```
## Loading required package: Matrix
```

```
## 
## Attaching package: 'Matrix'
```

```
## The following objects are masked from 'package:tidyr':
## 
##     expand, pack, unpack
```

```
## Warning: package 'lmerTest' was built under R version 4.5.2
```

```
## 
## Attaching package: 'lmerTest'
```

```
## The following object is masked from 'package:lme4':
## 
##     lmer
```

```
## The following object is masked from 'package:stats':
## 
##     step
```

```
## Warning: package 'performance' was built under R version 4.5.2
```

```
## 
## Attaching package: 'purrr'
```

```
## The following object is masked from 'package:caret':
## 
##     lift
```

```
## This is DHARMa 0.4.7. For overview type '?DHARMa'. For recent changes, type news(package = 'DHARMa')
```

```
## 
## Attaching package: 'nlme'
```

```
## The following object is masked from 'package:lme4':
## 
##     lmList
```

```
## The following object is masked from 'package:dplyr':
## 
##     collapse
```

```
## Loading required package: viridisLite
```

### Load and process data

```
#### Load data ####

##### load manually checked data (checked survival status manually) to see how censor/status aligns, includes both blocks! #####

fly_for<-read.csv("/Users/mkutzer/Desktop/Critical transitions/Data/Infected data/Infected_data_forward_checked.csv", header=TRUE)

#str(fly_for)

fly_for %>%
  dplyr::distinct(FlyID, Treatment) %>% # 440 Infected, 28 uninfected
  dplyr::count(Treatment)
```

```
##   Treatment   n
## 1  Infected 440
## 2        UI  28
```

```
# Set experiment end time (to truncate the data since both blocks ran longer than 14 days)
EXP_END <- 20160

# Discards all individuals that die before 10 hpi, selects 1:20160 minutes - this will need to be taken into account for the PE analysis!
# THIS IS THE ORIGINAL BLOCK
##### Block A #####
# 1. Block A: readRDS -> LONG (NO filtering yet)
setwd("/Users/mkutzer/Desktop/Critical transitions/R/")
rds_raw <- readRDS("data4ale.rds")

blockA_long <- rds_raw %>%
  # assumes 1st col is FlyID and cols 1:20160 exist as X1..X20160 or similar
  pivot_longer(cols = 1:20160, names_to = "time", values_to = "activity") %>%
  # keep only what we actually need!
  transmute(FlyID = as.character(FlyID), 
            time = as.numeric(gsub("\\D+", "", time)), # "X123" -> 123 (works for X-prefixed colnames) 
            activity = as.integer(activity), 
            source_block = "blockA_rds")

#### Block B: Kabir CSVs: LONG (NO filtering yet) ####

file_paths <- c("~/Desktop/Critical transitions/Data/Infected data/Data_PreProcessed_Kabir1CTM012_DA.csv", 
                "~/Desktop/Critical transitions/Data/Infected data/Data_PreProcessed_Kabir1CTM013_DA.csv", 
                "~/Desktop/Critical transitions/Data/Infected data/Data_PreProcessed_Kabir1CTM014_DA.csv", 
                "~/Desktop/Critical transitions/Data/Infected data/Data_PreProcessed_Kabir1CTM015_DA.csv", 
                "~/Desktop/Critical transitions/Data/Infected data/Data_PreProcessed_Kabir1CTM016_DA.csv", 
                "~/Desktop/Critical transitions/Data/Infected data/Data_PreProcessed_Kabir1CTM017_DA.csv", 
                "~/Desktop/Critical transitions/Data/Infected data/Data_PreProcessed_Kabir1CTM018_DA.csv", 
                "~/Desktop/Critical transitions/Data/Infected data/Data_PreProcessed_Kabir1CTM019_DA.csv")

blockB_long <- purrr::map_dfr(file_paths, function(fp) {
  wide <- readr::read_csv(fp, show_col_types = FALSE)
  
  # If these CSVs have a time column, drop it here
  # If they do NOT, this will just keep all cols as fly columns.
  # Adjust "select" as needed
  wide2 <- wide %>%
    select(-any_of(c("time","Time","minute","Minute","minutes","Minutes")))
  
  wide2 %>%
    mutate(time = row_number()) %>%
    pivot_longer(cols = -time, names_to = "FlyID", values_to = "activity") %>%
    transmute(FlyID = as.character(FlyID), 
              time = as.numeric(time), 
              activity = as.integer(activity), source_block = basename(fp))})

##### Combine both blocks of data #####
fly_raw <- bind_rows(blockA_long, blockB_long) %>% 
  filter(time >= 1, time <= EXP_END)

fly_raw %>% distinct(FlyID) %>% summarise(n_flies = n())
```

```
## # A tibble: 1 × 1
##   n_flies
##     <int>
## 1     468
```

```
##### Match FlyIDs to fly_data_pre and look at status/censor/class for each #####
# reference status from fly_for
fly_for_ref <- fly_for %>%
  transmute(FlyID = as.character(FlyID), 
            status_for = if_else(Status == 1, "dead", "alive"), 
            Treatment_for = as.character(Treatment), 
            LastMovementTime_for = as.numeric(LastMovementTime)) %>% distinct(FlyID, .keep_all = TRUE)

#### Join TRUE labels ONCE, then filter/censor 1x only! ####

# Join checked metadata onto raw minute-level activity
tmp <- fly_raw %>% 
  left_join(fly_for_ref, by = "FlyID")

# Identify FlyIDs that failed to join (or have incomplete metadata)
missing_ids <- tmp %>%
  filter(is.na(status_for) | is.na(Treatment_for) | is.na(LastMovementTime_for)) %>%
  distinct(FlyID)

if (nrow(missing_ids) > 0) {
  print(missing_ids, n = Inf)  # show all missing IDs
  stop("FlyIDs missing from Infected_data_forward_checked.csv (or have missing fields).")}

# Apply censoring and class definitions 1x
fly_data_pre <- tmp %>%
  mutate(Status = status_for, 
         Treatment = Treatment_for,
         LastMovementTime = LastMovementTime_for,
    
    # alive scored to EXP_END (experiment end); dead scored to LastMovementTime
    AnchorTime = ifelse(Status == "alive", EXP_END, LastMovementTime),
    
    class = case_when(Status == "dead"  & LastMovementTime < 600 ~ "discard", 
                      Status == "dead"  & LastMovementTime >= 600 ~ "dead", 
                      Status == "alive" ~ "alive")) %>%
  
  filter(class != "discard") %>%
  group_by(FlyID) %>%
  filter(time <= AnchorTime) %>%
  ungroup() %>%
  select(FlyID, Status, Treatment, time, activity, LastMovementTime, AnchorTime, class, LastMovementTime_for)

# drop this fly manually
fly_data_pre <- fly_data_pre %>%
  filter(FlyID != "M16T25_P4Dead")

# override checked LMT for specific FlyIDs *before* censoring
# Only do this if you have a correction to fly_for itself.
lmt_fix <- tibble(FlyID = c("M15T21_P4Dead"), 
                          LastMovementTime_for = c(1761))

fly_data_pre <- fly_data_pre %>%
  left_join(lmt_fix, by = "FlyID", suffix = c("", "_fix")) %>%
  mutate(LastMovementTime_for = coalesce(LastMovementTime_for_fix, LastMovementTime_for),
    LastMovementTime = LastMovementTime_for, 
    AnchorTime = ifelse(Status == "alive", EXP_END, LastMovementTime)) %>% 
  select(-LastMovementTime_for_fix)

# Re-apply censoring after any LMT edits
fly_data_pre <- fly_data_pre %>%
  group_by(FlyID) %>%
  filter(time <= AnchorTime) %>%
  ungroup()

fly_data_pre%>% distinct(FlyID) %>% summarise(n_flies = n())
```

```
## # A tibble: 1 × 1
##   n_flies
##     <int>
## 1     437
```

```
##### AND NOW SPLIT THE CONTROLS AND INFECTED #####
# drop old class (I checked everything and don't need it!), old_class is no longer included in this code
#fly_data_pre <- dplyr::select(fly_data_pre, -class_old) 

infected_df  <- dplyr::filter(fly_data_pre, Treatment == "Infected")
infected_df%>% distinct(FlyID) %>% summarise(n_flies = n())
```

```
## # A tibble: 1 × 1
##   n_flies
##     <int>
## 1     411
```

```
# View last movement/ time of death
infected_df %>% 
  distinct(FlyID, class, LastMovementTime) %>%
  arrange(class, LastMovementTime) %>% print(n=10)
```

```
## # A tibble: 411 × 3
##    FlyID          class LastMovementTime
##    <chr>          <chr>            <dbl>
##  1 M16T2_G3Dead   alive            20148
##  2 M12T12_P3Dead  alive            20151
##  3 M15T26_C1Dead  alive            20158
##  4 M12T3_G1Dead   alive            20160
##  5 M12T4_P1Dead   alive            20160
##  6 M12T7_G2Dead   alive            20160
##  7 M12T10_W3Alive alive            20160
##  8 M12T15_G4Alive alive            20160
##  9 M12T16_P4Alive alive            20160
## 10 M12T17_C4Dead  alive            20160
## # ℹ 401 more rows
```

```
fly_data <- dplyr::filter(fly_data_pre, Treatment == "Infected") # same as infected_df but now previous code runs
control_df <- dplyr::filter(fly_data_pre, Treatment == "UI")

##### Now proceed with processing but filter out uninfected individuals... #####
# continue with processing steps from above

# keep only infected flies before splitting
infected_list <- fly_data_pre %>%
  filter(Treatment != "UI") %>% # remove UIs
  split(.$FlyID)

# controls in their own list
control_list <- fly_data_pre %>%
  filter(Treatment == "UI") %>%
  split(.$FlyID)

###### Now fly_data only includes infected flies ######
fly_data = lapply(infected_list, function(df) {
  df = as.data.frame(df) 
  colnames(df)[colnames(df) == "FlyID"] = "ID"
  colnames(df)[colnames(df) == "time"] = "x"
  colnames(df)[colnames(df) == "activity"] = "y"
  return(df)})
```

### 1. Activity time series and figures

```
#### Figure 1 ####

# changed the code, previous code was grouping surviving flies as dead; also the code did not differentiate between uninfected and infected flies!

fly_data_hourly <- infected_df %>%
  select(FlyID, class, time, activity, LastMovementTime) %>%
  mutate(hour = floor(time / 60), 
         deadhour = floor(LastMovementTime / 60)) %>%
  group_by(FlyID, class, hour) %>%     # <- remove deadhour from grouping
  summarise(activity = mean(activity, na.rm = TRUE), 
            deadhour = first(deadhour), # keep it for plotting/annotations 
            .groups = "drop")


fly_data_hourly %>% distinct(FlyID) %>% summarise(n_flies = n())
```

```
## # A tibble: 1 × 1
##   n_flies
##     <int>
## 1     411
```

```
mean_data_hourly <- fly_data_hourly %>%
  group_by(class, hour) %>%  
  summarise(mean_activity = mean(activity, na.rm = TRUE), 
            .groups = "drop")

#ploties.1 = c() # this seems superfluous and overly complicated... maybe I missed something?

# One row per fly with its hour of death
deaths <- infected_df %>%
  distinct(FlyID, Treatment, class, LastMovementTime) %>% 
  filter(class == "dead") %>%
  mutate(death_hour = floor(LastMovementTime/ 60))

# Count deaths per hour (per treatment)
deaths_per_hour <- deaths %>%
  group_by(Treatment, death_hour) %>%
  summarise(n_deaths = n(), .groups = "drop")

###### Plot deaths per hour ######
deaths_hour <- ggplot(deaths_per_hour, aes(x = death_hour, y = n_deaths)) +
  geom_col(color = "black", fill = "#D55E00") +
  #facet_wrap(~ Treatment, scales = "free_y") +
  scale_x_continuous(
    breaks = seq(0, max(deaths_per_hour$death_hour), by = 25)) +
  labs(x = "Hour", y = "# deaths per hour") +
  theme_minimal(); deaths_hour
```

```
# same with 2 hour bins (previous plot was in 3.3. hour bins for some reason)
deaths_per_hour_binned <- deaths %>%
  mutate(death_hour_bin = floor(death_hour / 2) * 2) %>% # e.g. 0–1 is 0, 2–3 is 2
  group_by(Treatment, death_hour_bin) %>%
  summarise(n_deaths = n(), .groups = "drop")

###### Plot deaths - 2 hour bins ######
deaths_binned <- ggplot(deaths_per_hour_binned, aes(x = death_hour_bin, y = n_deaths)) +
  geom_col(color = "black", fill = "#D55E00", width = 2) +   # width = 2 to match bins
  #facet_wrap(~ Treatment, scales = "free_y") +
  scale_x_continuous(
    breaks = seq(0, max(deaths_per_hour_binned$death_hour_bin), by = 25)) +
  labs(x = "Hours post infection", y = "# deaths") +
  theme_minimal() +
  theme(axis.text = element_text(size=14),
        axis.title = element_text(size = 16)); deaths_binned
```

```
###### Activity time series ######
activity_time_series <- ggplot() +
  geom_line(data = fly_data_hourly, aes(x = hour, y = activity, group = FlyID, color = class), alpha = 0.1) +
  geom_line(data = mean_data_hourly, aes(x = hour, y = mean_activity), size = 1.5, color = "gray4") +
  facet_wrap(~class, ncol = 1) +  # Separate panels for each class
  scale_x_continuous(expand = c(0,0), breaks = seq(0, max(fly_data_hourly$hour, na.rm = TRUE), by = 25)) +
  labs(x = "Hours post infection",
       y = "Mean activity",
       color = "Class") +
  theme_minimal() +
  theme(legend.position = "none",
        axis.text = element_text(size=14),
        axis.title = element_text(size = 16))+
  scale_color_manual(values = c("alive" = "#0072B2", "dead" = "#D55E00")); activity_time_series
```

```
## Warning: Using `size` aesthetic for lines was deprecated in ggplot2 3.4.0.
## ℹ Please use `linewidth` instead.
## This warning is displayed once every 8 hours.
## Call `lifecycle::last_lifecycle_warnings()` to see where this warning was
## generated.
```

```
fly_data_hourly %>%
  dplyr::distinct(FlyID, class) %>%   # one row per fly
  dplyr::count(class, name = "n_flies")
```

```
## # A tibble: 2 × 2
##   class n_flies
##   <chr>   <int>
## 1 alive     161
## 2 dead      250
```

```
###### Create Figure 1 using plot_grid ######
Fig1 <- plot_grid(deaths_binned, activity_time_series, 
                  labels = c("A", "B"), rel_widths = c(1,1), rel_heights = c(1,2),
                  label_size = 18, ncol = 1, align = "hv", axis = c("lr"))
Fig1
```

```
save_plot("/Users/mkutzer/Desktop/Critical transitions/Manuscript/Figure_1.png", base_width = 10, base_height = 6, Fig1)
```

### 1a. Uninfected time series and deaths/ 2hrs

```
#### Create an uninfected df ####
uninfected_df  <- dplyr::filter(fly_data_pre, Treatment == "UI")
uninfected_df%>% distinct(FlyID) %>% summarise(n_flies = n())
```

```
## # A tibble: 1 × 1
##   n_flies
##     <int>
## 1      26
```

```
ui_fly_data_hourly <- uninfected_df %>%
  select(FlyID, class, time, activity, LastMovementTime) %>%
  mutate(hour = floor(time/ 60), 
         deadhour = floor(LastMovementTime/ 60)) %>%
  group_by(FlyID, class, hour) %>%     # <- remove deadhour from grouping
  summarise(activity = mean(activity, na.rm = TRUE), 
            deadhour = first(deadhour), # keep for plotting/annotations 
            .groups = "drop")


ui_fly_data_hourly %>%
  dplyr::distinct(FlyID, class) %>%   # one row per fly
  dplyr::count(class, name = "n_flies")
```

```
## # A tibble: 2 × 2
##   class n_flies
##   <chr>   <int>
## 1 alive      22
## 2 dead        4
```

```
ui_mean_data_hourly <- ui_fly_data_hourly %>%
  group_by(class, hour) %>%  
  summarise(
    mean_activity = mean(activity, na.rm = TRUE),
    .groups = "drop")

#### deaths per 2-hour bin (UI) ####
ui_deaths <- uninfected_df %>%
  distinct(FlyID, class, LastMovementTime) %>%
  filter(class == "dead") %>%
  mutate(death_hour = floor(LastMovementTime/ 60),
         death_hour_bin = floor(death_hour/ 2) * 2)

ui_deaths_per_bin <- ui_deaths %>%
  count(death_hour_bin, name = "n_deaths")

ui_deaths_binned <- ggplot(ui_deaths_per_bin, aes(x = death_hour_bin, y = n_deaths)) +
  geom_col(color = "black", fill = "#D55E00", width = 2) +
  scale_y_continuous(limits = c(0,35)) +
  scale_x_continuous(breaks = seq(0, max(ui_deaths_per_bin$death_hour_bin, na.rm = TRUE), by = 25)) +
  labs(x = "Hours post infection", y = "# deaths") +
  theme_minimal() +
  theme(axis.text = element_text(size=14),
        axis.title = element_text(size = 16))

ui_deaths_binned
```

```
ui_activity_time_series <- ggplot() +
  geom_line(data = ui_fly_data_hourly, aes(x = hour, y = activity, group = FlyID, color = class), alpha = 0.5) +
  geom_line(data = ui_mean_data_hourly, aes(x = hour, y = mean_activity), size = 1.5, color = "gray4") +
  facet_wrap(~class, ncol = 1) +  # Separate panels for each class
  scale_x_continuous(expand = c(0,0), breaks = seq(0, max(ui_fly_data_hourly$hour, na.rm = TRUE), by = 25)) +
  scale_y_continuous(limits = c(0, 15)) +
  labs(x = "Hours post infection",
       y = "Mean activity",
       color = "Class") +
  theme_minimal() +
  theme(legend.position = "none",
        axis.text = element_text(size=14),
        axis.title = element_text(size = 16))+
  scale_color_manual(values = c("alive" = "#0072B2", "dead" = "#D55E00")); ui_activity_time_series
```

```
Fig_UI <- plot_grid(ui_deaths_binned, ui_activity_time_series,
  labels = c("A", "B"),
  rel_heights = c(1, 2),
  label_size = 18,
  ncol = 1, align = "hv", axis = "lr")

#### Figure S1 ####
Fig_UI
```

```
#### Save uninfected figure as Figure S1 ####
save_plot("/Users/mkutzer/Desktop/Critical transitions/Manuscript/Figure_S1.png", base_width = 10, base_height = 6, Fig_UI)

summ14 <- ui_fly_data_hourly %>% 
  group_by(FlyID, class) %>%       
  summarise(n_hours = sum(!is.na(activity)), 
            mean14 = mean(activity, na.rm = TRUE), 
            sd14 = sd(activity, na.rm = TRUE), 
            prop0_14 = mean(activity == 0, na.rm = TRUE), 
            .groups = "drop")
summ14
```

```
## # A tibble: 26 × 6
##    FlyID               class n_hours mean14  sd14 prop0_14
##    <chr>               <chr>   <int>  <dbl> <dbl>    <dbl>
##  1 M13T10_ControlDead  alive     337   1.14  1.37   0.181 
##  2 M13T11_ControlDead  alive     337   1.68  1.60   0.107 
##  3 M14T13_ControlAlive alive     337   3.95  3.07   0.0890
##  4 M14T27_ControlDead  alive     337   2.51  2.32   0.0920
##  5 M14T28_ControlDead  dead       78   3.38  2.01   0.0641
##  6 M14T29_ControlAlive alive     337   1.69  1.54   0.0920
##  7 M15T31_ControlDead  alive     337   2.11  1.79   0.0979
##  8 M15T32_ControlDead  alive     337   1.30  1.76   0.145 
##  9 M15T6_ControlDead   alive     337   1.87  2.33   0.220 
## 10 M16T10_ControlAlive alive     337   1.88  1.96   0.131 
## # ℹ 16 more rows
```

```
group_means14 <- summ14 %>%
  dplyr::group_by(class) %>%
  dplyr::summarise(
    n_flies = dplyr::n(),
    mean_of_fly_means = mean(mean14, na.rm = TRUE),
    sd_of_fly_means   = sd(mean14, na.rm = TRUE),
    se = sd_of_fly_means / sqrt(n_flies),
    .groups = "drop")
group_means14
```

```
## # A tibble: 2 × 5
##   class n_flies mean_of_fly_means sd_of_fly_means    se
##   <chr>   <int>             <dbl>           <dbl> <dbl>
## 1 alive      22              2.21           0.719 0.153
## 2 dead        4              3.57           1.42  0.709
```

```
summ14_i <- fly_data_hourly %>%
  group_by(FlyID, class) %>%       
  summarise(
    n_hours = sum(!is.na(activity)),
    mean14  = mean(activity, na.rm = TRUE),
    sd14    = sd(activity, na.rm = TRUE),
    prop0_14 = mean(activity == 0, na.rm = TRUE),
    .groups = "drop")
summ14_i
```

```
## # A tibble: 411 × 6
##    FlyID       class n_hours mean14  sd14 prop0_14
##    <chr>       <chr>   <int>  <dbl> <dbl>    <dbl>
##  1 AliveM12T11 alive     337   2.12  1.85   0.0801
##  2 AliveM12T19 alive     337   1.03  1.36   0.237 
##  3 AliveM12T26 alive     337   1.78  1.71   0.142 
##  4 AliveM12T27 alive     337   5.10  4.11   0.134 
##  5 AliveM12T28 alive     337   3.00  2.35   0.0950
##  6 AliveM12T5  alive     337   2.76  1.93   0.0682
##  7 AliveM12T8  alive     337   2.32  2.23   0.0890
##  8 AliveM13T18 alive     337   2.94  1.94   0.0208
##  9 AliveM13T6  alive     337   2.04  1.99   0.154 
## 10 AliveM14T22 alive     337   2.50  1.96   0.0890
## # ℹ 401 more rows
```

```
group_means14_i <- summ14_i %>%
  dplyr::group_by(class) %>%
  dplyr::summarise(
    n_flies = dplyr::n(),
    mean_of_fly_means = mean(mean14, na.rm = TRUE),
    sd_of_fly_means   = sd(mean14, na.rm = TRUE),
    se = sd_of_fly_means / sqrt(n_flies),
    .groups = "drop")
group_means14_i
```

```
## # A tibble: 2 × 5
##   class n_flies mean_of_fly_means sd_of_fly_means     se
##   <chr>   <int>             <dbl>           <dbl>  <dbl>
## 1 alive     161              2.18           0.744 0.0587
## 2 dead      250              2.93           1.80  0.114
```

### 2. Early warning metrics and analyses

Uninfected controls removed

```
#### Figure 2 ####

##### run_dynfoot #####
fly_output <- run_dynfoot(
  fly_data,
  metrics.type = "all",
  detrend.type = "none",
  group = "class",
  cores = 1,
  saveplots = FALSE
)

##### Define metrics #####
metrics <- c("mean.full", "SD.full","ar1.full", "CV.full",
             "mean.roll", "SD.roll","ar1.roll", "CV.roll")
group <- "class"
full_metrics <- metrics[stringr::str_detect(metrics, "full")]
roll_metrics <- metrics[stringr::str_detect(metrics, "roll")]

##### Build EWS summary table #####
ews_df <- sort_ews(do.call(rbind, fly_output), group, metrics)

stopifnot(all(metrics %in% names(ews_df)))
stopifnot("ID" %in% names(ews_df))
stopifnot(group %in% names(ews_df))  # "class" exists here

##### Rank-normalise FULL metrics only #####
ews_df[full_metrics] <- lapply(ews_df[full_metrics], norm_ts, type = "rank")

##### Rename group column to 'clase' (this was a mistake in the original analysis and I just kept it to avoid confusion) #####
names(ews_df)[names(ews_df) == group] <- "clase"

##### Add block and modelling factors #####
df2 <- ews_df %>%
  dplyr::mutate(block = dplyr::case_when(
    stringr::str_detect(ID, "^(Alive|Dead)") ~ "Block1", 
    stringr::str_detect(ID, "^M\\d+") ~ "Block2",
    TRUE ~ NA_character_),
    block = factor(block),
    clase_f = factor(clase),
    FlyID = ID)

stopifnot(!anyNA(df2$block))

##### Statistical Modelling: Fit one LMM per metric and extract Type II main-effect p-values #####
long <- df2 %>%
  tidyr::pivot_longer(all_of(metrics), names_to = "metric", values_to = "value") %>%
  dplyr::filter(!is.na(value)) %>%
  dplyr::mutate(metric = factor(metric, levels = metrics))

models_by_metric <- long %>%
  dplyr::group_by(metric) %>%
  dplyr::group_map(~ lmerTest::lmer(value ~ clase_f + (1 | block), data = .x, REML = TRUE))
```

```
## boundary (singular) fit: see help('isSingular')
## boundary (singular) fit: see help('isSingular')
```

```
names(models_by_metric) <- levels(long$metric)

get_p <- function(a, term) {
  if (!term %in% rownames(a)) return(NA_real_)
  if ("Pr(>Chisq)" %in% colnames(a)) return(a[term, "Pr(>Chisq)"])
  if ("Pr(>F)"  %in% colnames(a)) return(a[term, "Pr(>F)"])
  NA_real_}

##### Store the statistics! #####
anova_main <- purrr::imap_dfr(models_by_metric, ~{a <- car::Anova(.x, type = 2) # car Anova for main effects
  tibble::tibble(metric = .y, p_class_main = get_p(a, "clase_f"))}) %>%
  dplyr::mutate(p_class_main_BH = p.adjust(p_class_main, method = "BH")) # Benjamini and Hochberg corrected p-values

##### Add stars from BH adjusted p-values for figures #####
alpha_stars <- function(q){
  dplyr::case_when(
    is.na(q) ~ "",
    q < 0.001 ~ "***",
    q < 0.01 ~ "**",
    q < 0.05 ~ "*",
    q < 0.1 ~ "·",  
    TRUE ~ "ns")}

label_map <- anova_main %>%
  dplyr::transmute(metric, q = p_class_main_BH, stars = alpha_stars(q))

##### FULL metrics plot #####
metric_labs_full <- c("mean.full" = "Mean",
                      "SD.full" = "SD",
                      "ar1.full" = "Lag-1 AC",
                      "CV.full" = "CV")

temp_full <- as.data.frame(df2) %>%
  dplyr::add_count(clase) %>%
  dplyr::mutate(clase = paste0(clase, " (", n, ")")) %>%
  reshape2::melt(measure.vars = full_metrics, variable.name = "metric") %>%
  dplyr::left_join(label_map, by = "metric")

full_metric_plot <- ggplot(temp_full, aes(x = metric, y = value, fill = clase)) +
geom_boxplot(outlier.shape = NA, alpha = 0.8, na.rm = TRUE, 
             position = ggplot2::position_dodge()) +
geom_point(na.rm = TRUE, position = ggplot2::position_jitterdodge(0.07), pch = 21) + 
  labs(x = "Metric", y = "Percentile rank") + 
  scale_x_discrete(labels = metric_labs_full) +
  theme_light() +
  scale_fill_manual(values = c("#0072B2", "#D55E00")) +
  theme(axis.text = element_text(size = 14),
                 axis.title = element_text(size = 16),
                 legend.text = element_text(size = 12),
                 legend.title = element_text(size = 14)) +
  geom_text(data = distinct(temp_full, metric, stars), aes(x = metric, y = 1.05, label = stars),
            inherit.aes = FALSE, fontface = "bold") + 
  labs(fill = "Class (# obs)")

##### ROLLING metrics plot #####
metric_labs_roll <- c("mean.roll" = "Mean",
                      "SD.roll" = "SD",
                      "ar1.roll" = "Lag-1 AC",
                      "CV.roll" = "CV")

temp_roll <- as.data.frame(df2) %>%
  dplyr::add_count(clase) %>%
  dplyr::mutate(clase = paste0(clase, " (", n, ")")) %>%
  reshape2::melt(measure.vars = roll_metrics, variable.name = "metric") %>%
  dplyr::left_join(label_map, by = "metric")

roll_ypos <- temp_roll %>%
  dplyr::group_by(metric) %>%
  dplyr::summarise(y_star = max(value, na.rm = TRUE) + 0.08,
                   stars = dplyr::first(stars),
                   .groups = "drop")

rolling_metric_plot <- ggplot(temp_roll, ggplot2::aes(x = metric, y = value, fill = clase)) +
 geom_boxplot(outlier.shape = NA, alpha = 0.8, na.rm = TRUE,
                        position = position_dodge()) +
  geom_point(na.rm = TRUE, position = position_jitterdodge(0.05), pch = 21) +
  labs(x = "Metric", y = "Kendall's tau") +
  scale_x_discrete(labels = metric_labs_roll) +
  theme_light() +
  theme(axis.text = element_text(size = 14),
                 axis.title = element_text(size = 16),
                 legend.text = element_text(size = 12),
                 legend.title = element_text(size = 14)) +
  scale_fill_manual(values = c("#0072B2", "#D55E00")) +
  geom_text(data = roll_ypos,
                     ggplot2::aes(x = metric, y = y_star, label = stars),
                     inherit.aes = FALSE, fontface = "bold") + 
  labs(fill = "Class (# obs)"); rolling_metric_plot
```

```
##### Combine plots and save Figure 2 #####
legend <- cowplot::get_legend(rolling_metric_plot + ggplot2::theme(legend.position = "right"))

full_metric_plot_clean <- full_metric_plot + ggplot2::theme(legend.position = "none")
rolling_metric_plot_clean <- rolling_metric_plot + ggplot2::theme(legend.position = "none")

plots <- plot_grid(full_metric_plot_clean, rolling_metric_plot_clean,
                            nrow = 2, labels = c("A", "B"))

Figure_2 <- plot_grid(plots, legend, ncol = 2, rel_widths = c(1, 0.2))

Figure_2
```

```
save_plot("/Users/mkutzer/Desktop/Critical transitions/Manuscript/Figure_2.png", base_width = 8, base_height = 6, Figure_2)

#### STATISTICS: rolling and full ####
anova_main
```

```
## # A tibble: 8 × 3
##   metric    p_class_main p_class_main_BH
##   <chr>            <dbl>           <dbl>
## 1 mean.full   0.0000732       0.000293  
## 2 SD.full     0.00000122      0.00000972
## 3 ar1.full    0.00619         0.0103    
## 4 CV.full     0.0292          0.0390    
## 5 mean.roll   0.254           0.254     
## 6 SD.roll     0.000658        0.00176   
## 7 ar1.roll    0.0587          0.0671    
## 8 CV.roll     0.00641         0.0103
```

### 3. Random forests

```
#### Random forest models ####
# Before sort_ews: all fly IDs in fly_output
all_ids <- unique(do.call(rbind, fly_output)$ID)
length(all_ids)
```

```
## [1] 411
```

```
full.data = sort_ews(data = do.call(rbind,fly_output), group = "class",metrics = metrics)
nrow(full.data)
```

```
## [1] 411
```

```
colnames(full.data)[colnames(full.data) == "class"] = "Status"

set.seed(999)
train_indices = sample(1:nrow(full.data), size = 0.6*nrow(full.data)) # sample 60% data for training, these are randomly sampled - Measurements are from 1 fly each! 
train_data = full.data[train_indices, ] 
train_data$Status = as.factor(train_data$Status)
test_data = full.data[-train_indices, ] 

# Convert 'Status' from numeric to factor in training and test data 
train_data$Status = as.factor(train_data$Status)
test_data$Status = as.factor(test_data$Status)
train_data$Status = factor(train_data$Status, levels = c("dead", "alive"))
test_data$Status = factor(test_data$Status, levels = c("dead", "alive"))
head(train_data)
```

```
## # A tibble: 6 × 12
##   ID        Status ts.length LastMovementTime mean.full SD.full ar1.full CV.full
##   <chr>     <fct>      <int>            <dbl>     <dbl>   <dbl>    <dbl>   <dbl>
## 1 M19T9_W1… dead        1618             1618     2.61     4.23    0.456    1.62
## 2 M16T28_A… dead       17452            17452     0.447    2.07    0.828    4.62
## 3 DeadM13T… dead        1009             1009     2.66     4.84    0.399    1.82
## 4 DeadM14T1 dead        1842             1842     3.60     5.94    0.636    1.65
## 5 M17T9_W3… dead        1801             1801     4.22     4.35    0.628    1.03
## 6 DeadM17T… alive      20160            20160     2.42     3.80    0.622    1.57
## # ℹ 4 more variables: mean.roll <dbl>, SD.roll <dbl>, ar1.roll <dbl>,
## #   CV.roll <dbl>
```

```
set.seed(999)

##### Run the model, LEAVE IN WINDOWS #####
rf_model = randomForest(Status ~ .- ID - ts.length -LastMovementTime, data = train_data, ntree = 300, importance = TRUE) # train the model
test_pred = predict(rf_model, newdata = test_data) # test the model

##### Confusion Matrix #####
cm_full = confusionMatrix(test_pred, test_data$Status) 
cm_full
```

```
## Confusion Matrix and Statistics
## 
##           Reference
## Prediction dead alive
##      dead    92    13
##      alive    7    53
##                                          
##                Accuracy : 0.8788         
##                  95% CI : (0.819, 0.9244)
##     No Information Rate : 0.6            
##     P-Value [Acc > NIR] : 2.594e-15      
##                                          
##                   Kappa : 0.7436         
##                                          
##  Mcnemar's Test P-Value : 0.2636         
##                                          
##             Sensitivity : 0.9293         
##             Specificity : 0.8030         
##          Pos Pred Value : 0.8762         
##          Neg Pred Value : 0.8833         
##              Prevalence : 0.6000         
##          Detection Rate : 0.5576         
##    Detection Prevalence : 0.6364         
##       Balanced Accuracy : 0.8662         
##                                          
##        'Positive' Class : dead           
##
```

```
##### Store cm in a df #####
cm_df <- as.data.frame(cm_full$table)
groups = c("dead","alive")

# Normalize frequencies within each row (actual class) to calculate relative frequencies
cm_df <- cm_df %>%
  group_by(Reference) %>%
  mutate(RelativeFreq = Freq/ sum(Freq))

# Create a discrete color palette for each class
n_cols <- length(unique(cm_df$Reference))  # Number of unique classes
viridis_palette <- viridis(n_cols, option = "H")  # Generate Viridis colors, not used

our_palette = c("dead" = "#D55E00", "alive" = "#0072B2")

# Assign colors to each class based on column order
cm_df <- cm_df %>%
  dplyr::mutate(Color = our_palette[Reference])

# Generate the confusion matrix plot using ggplot2
cm_full_plot <- ggplot(cm_df, aes(x = Reference, y = Prediction)) +
  geom_tile(aes(fill = Color, alpha = RelativeFreq), color = "white") +
  geom_text(aes(label = sprintf("%1.0f", Freq))) +
  scale_fill_identity() +
  scale_alpha(range = c(0.5, 1)) +
  labs(x = "actual", y = "predicted") +
  theme_minimal() +
  theme(
    legend.position = "none",
    panel.grid = element_blank(),
    plot.margin = ggplot2::margin(2, 2, 2, 2)); cm_full_plot
```

```
##### Plot of variable importance: full model #####
varImpPlot(rf_model)
```

```
# Predict probabilities
prob_predictions = predict(rf_model, newdata = test_data, type = "prob")
#prob_predictions (view if needed)

# Extract probabilities for the class 'dead' which represents death
death_probs = prob_predictions[, "dead"]

# Convert the test data 'Status' from a factor to numeric values
numeric_status = ifelse(test_data$Status=="dead",1,0)

##### Generate the ROC curve using the pROC package #####
roc_obj = roc(numeric_status, death_probs)
```

```
## Setting levels: control = 0, case = 1
```

```
## Setting direction: controls < cases
```

```
###### AUC ######
auc = roc_obj$auc # 0.8584

# Convert ROC object to data frame
roc_df = data.frame(
  FPR = 1 - roc_obj$specificities,
  TPR = roc_obj$sensitivities)

###### Plot the roc curve ######
roc_curve_full <- ggplot(roc_df, aes(x = FPR, y = TPR)) + 
  geom_path(linewidth = 1, color = "#0072B2") + 
  geom_abline(slope = 1, intercept = 0, linetype = "dashed", color = "gray") +
  labs(x = "False positive rate", y = "True positive rate") + 
  theme_minimal() +
  coord_cartesian(xlim = c(0, 1), ylim = c(0, 1), expand = FALSE) +
  annotate("text", x = 0.25, y = 0.65,
           label = paste0("AUC = ", round(as.numeric(auc), 3)),
           hjust = 0)

###### Combine the confusion matrix and the roc curve ######

# adjust margins as needed on the cm
cm_inset_bordered <- ggdraw(cm_full_plot) +
  draw_grob(
    grid::rectGrob(gp = grid::gpar(fill = NA, col = "black", lwd = 0.8)))

fig_full_rf <- ggdraw(roc_curve_full) +
  draw_plot(cm_inset_bordered, x = 0.47, y = 0.15, width = 0.46, height = 0.46)

fig_full_rf
```

```
#### Random Forest: Windows #### 
#N.B! our_palette = c("dead" = "#D55E00", "alive" = "#0072B2")
auc = c()
start = c(240,480,720,960,1200,1440,1680,1920)
end = c(0,240,480,720,960,1200,1440,1680)

# inspect
roc_list <- vector("list", length(end))
cm_list  <- vector("list", length(end))

for (i in 1:length(end)) {
  
  fly_data = split(infected_df,infected_df$FlyID)
  
  fly_data = lapply(fly_data, function(df) {
    df = as.data.frame(df) %>% filter(time>LastMovementTime-start[i],time<LastMovementTime-end[i])
    colnames(df)[colnames(df) == "FlyID"] = "ID"
    colnames(df)[colnames(df) == "time"] = "x"
    colnames(df)[colnames(df) == "activity"] = "y"
    if (length(df$y)==0) {
      return(NULL)  # Exclude this data frame
    }
    return(df)
  })
  
  fly_data = fly_data[!sapply(fly_data, is.null)]
  
  fly_output = run_dynfoot_h(fly_data,metrics.type = "all",detrend.type = "none",group = "class",cores = if (.Platform$OS.type == "windows") 1 else 4, saveplots = FALSE)
  
  full.data = sort_ews(data = do.call(rbind,fly_output),group = "class",metrics = metrics[str_detect(metrics, "full")])
  
  full.data <- na.omit(full.data)
  
  set.seed(123)
  train_indices = sample(1:nrow(full.data), size = 0.6*nrow(full.data)) # sample 60 % data for training, these are randomly sampled - Measurements are from 1 fly each! 
  colnames(full.data)[colnames(full.data) == "class"] = "Status"
  train_data = full.data[train_indices, ] 
  train_data$Status = as.factor(train_data$Status)
  test_data = full.data[-train_indices, ]
  
  # Convert 'Status' from numeric to factor in training and test data 
  train_data$Status = as.factor(train_data$Status)
  test_data$Status = as.factor(test_data$Status)
  train_data$Status = factor(train_data$Status, levels = c("dead", "alive"))
  test_data$Status = factor(test_data$Status, levels = c("dead", "alive"))
  
  ## Get the nb of trees for OBB stabilising to a low rate
  set.seed(123)
  rf_modelWin = randomForest(Status ~ .- ID - ts.length - LastMovementTime, data = train_data, ntree = 300, importance = TRUE) # train the model
  test_pred = predict(rf_modelWin, newdata = test_data) # test the model
  
  cm_Win = confusionMatrix(test_pred, test_data$Status) # cm read in different way
  #print(cm_Win)
  
  # Plot of variable importance
  varImpPlot(rf_modelWin)
  
  # Predict probabilities
  prob_predictionsWin = predict(rf_modelWin, newdata = test_data, type = "prob")
  #prob_predictionsWin
  
  # Extract probabilities for the class 'dead' which represents death
  death_probsWin = prob_predictionsWin[, "dead"]
  
  # Convert the test data 'Status' from a factor to numeric values
  numeric_status = ifelse(test_data$Status=="dead",1,0)
  
  # Generate the ROC curve using the pROC package ##
  roc_objWin = roc(numeric_status, death_probsWin)
  
  auc[i] = roc_objWin$auc[1] # AUC  = area under the curve
  
  # store per-window outputs
  cm_list[[i]]  <- cm_Win
  roc_list[[i]] <- roc_objWin
}
```

```
## Setting levels: control = 0, case = 1
## Setting direction: controls < cases
```

```
## Setting levels: control = 0, case = 1
## Setting direction: controls < cases
```

```
## Setting levels: control = 0, case = 1
## Setting direction: controls < cases
```

```
## Setting levels: control = 0, case = 1
## Setting direction: controls < cases
```

```
## Setting levels: control = 0, case = 1
## Setting direction: controls < cases
```

```
## Setting levels: control = 0, case = 1
## Setting direction: controls < cases
```

```
## Setting levels: control = 0, case = 1
## Setting direction: controls < cases
```

```
## Setting levels: control = 0, case = 1
## Setting direction: controls < cases
```

```
# Choose which window to visualize
k <- which.max(auc)   

time <-  c("0-4", "4-8", "8-12", "12-16", "16-20", "20-24", "24-28", "28-32")
data <- data.frame(time = factor(time,levels=time), AUC = auc)

##### Create barplot with ggplot2 #####
AUC_windows <- ggplot(data, aes(x = time, y = AUC)) +
  geom_col(fill = "#D55E00") +
  scale_y_continuous(limits = c(0, 1), breaks = seq(0, 1, 0.2), expand = c(0, 0)) +
  labs(x = "Hours before death", y = "AUC") +
  theme_minimal() +
  theme(
    plot.margin = ggplot2::margin(5, 5, 5, 5))

# ROC for window k 
roc_k <- roc_list[[k]]
roc_df <- data.frame(FPR = 1 - roc_k$specificities, TPR = roc_k$sensitivities)

AUC_best <- ggplot(roc_df, aes(FPR, TPR)) +
  geom_path(linewidth=1, color = "#0072B2") +
  geom_abline(slope=1, intercept=0, linetype="dashed") +
  coord_equal() + theme_minimal() +
  labs(x="false positive rate", y="true positive rate") +
  annotate("text", x=0.25, y=0.35, label=paste0("AUC = ", round(auc[k], 2))); AUC_best
```

```
# Confusion matrix for window k 
cm_k <- as.data.frame(cm_list[[k]]$table)
cm_k <- cm_k %>%
  dplyr::group_by(Reference) %>%
  dplyr::mutate(RelativeFreq = Freq / sum(Freq)) %>%
  dplyr::ungroup() %>%
  dplyr::mutate(Color = c("alive"="#0072B2","dead"= "#D55E00")[as.character(Reference)])

cm_best_plot <- ggplot(cm_k, aes(x=Reference, y=Prediction)) +
  geom_tile(aes(fill=Color, alpha=RelativeFreq), color="white") +
  geom_text(aes(label=sprintf("%1.0f", Freq)), vjust=1) +
  scale_fill_identity() + scale_alpha(range=c(0.5,1)) +
  labs(x="actual", y="predicted", title=paste0("Window: ", time[k], " h")) +
  theme_minimal() + theme(legend.position="none", panel.grid=element_blank()); cm_best_plot
```

```
#### Combine plots for figure 4 ####

# combine
final_fig4 <- plot_grid(
  fig_full_rf,
  AUC_windows,
  labels = c("A", "B"),
  nrow = 1,
  align = "v",
  axis = "tblr",
  rel_widths = c(1, 1)
)

final_fig4
```

```
save_plot("/Users/mkutzer/Desktop/Critical transitions/Manuscript/Figure_4.png", base_width = 12, base_height = 6, final_fig4)
```

### 4. Permutation entropy

```
#### Parameters ####
bin_minutes <- 30L # PE window length (non-overlapping)
max_minutes_activity <- 20160L # 14 dpi used in activity pipeline
max_windows_14d <- max_minutes_activity/ bin_minutes  # 672
stopifnot(max_windows_14d == 672)

#### Load PE files ####

files <- c(
  "PermEnData_PreProcessed_Monitor 12 filtered.csv",
  "PermEnData_PreProcessed_Monitor 13 filtered.csv",
  "PermEnData_PreProcessed_Monitor 14 filtered.csv",
  "PermEnData_PreProcessed_Monitor 15 filtered.csv",
  "PermEnData_PreProcessed_Monitor 16 filtered.csv",
  "PermEnData_PreProcessed_Monitor 17 filtered.csv",
  "PermEnData_PreProcessed_Monitor 18 filtered.csv",
  "PermEnData_PreProcessed_Monitor 19 Filtered.csv",
  "PermEnData_PreProcessed_Kabir1CTM012_DA.csv",
  "PermEnData_PreProcessed_Kabir1CTM013_DA.csv",
  "PermEnData_PreProcessed_Kabir1CTM014_DA.csv",
  "PermEnData_PreProcessed_Kabir1CTM015_DA.csv", 
  "PermEnData_PreProcessed_Kabir1CTM016_DA.csv",
  "PermEnData_PreProcessed_Kabir1CTM017_DA.csv",
  "PermEnData_PreProcessed_Kabir1CTM018_DA.csv",
  "PermEnData_PreProcessed_Kabir1CTM019_DA.csv")

#### Read and reshape to long format (keep original IDs) ####
# drop _UnI (uninfected)
# create FlyID_key used for joining (same as cleaned ID)

pe_long <- purrr::map_dfr(files, function(f) {
  df <- readr::read_csv(f, show_col_types = FALSE)
  
  df %>%
    mutate(
      pe_window = dplyr::row_number(),
      source_file = f
    ) %>%
    tidyr::pivot_longer(
      cols = -c(pe_window, source_file),
      names_to = "FlyID_raw",
      values_to = "entropy"
    )
})

pe_long <- pe_long %>%
  mutate(
    FlyID_raw = stringr::str_trim(FlyID_raw),
    is_uninfected = stringr::str_detect(FlyID_raw, "_UnI$"),
    FlyID_key = stringr::str_replace(FlyID_raw, "_UnI$", "")  # only strip suffix
  ) %>%
  filter(!is_uninfected) %>%
  select(-is_uninfected)

truth_ids <- unique(stringr::str_trim(as.character(fly_for$FlyID)))

mean(pe_long$FlyID_key %in% truth_ids)
```

```
## [1] 0.9973321
```

```
anti <- pe_long %>%
  distinct(FlyID_raw, FlyID_key) %>%
  filter(!(FlyID_key %in% truth_ids))

print(head(anti, 50))
```

```
## # A tibble: 1 × 2
##   FlyID_raw  FlyID_key 
##   <chr>      <chr>     
## 1 DeadM19T25 DeadM19T25
```

```
#### Trim terminal zero padding (per fly) ####

trim_terminal_zero_padding <- function(df) {
  df %>%
    group_by(FlyID_key) %>%
    mutate(
      last_nonzero_window = suppressWarnings(max(pe_window[!is.na(entropy) & entropy != 0], na.rm = TRUE))
    ) %>%
    filter(is.finite(last_nonzero_window), pe_window <= last_nonzero_window) %>%
    select(-last_nonzero_window) %>%
    ungroup()
}

pe_long <- trim_terminal_zero_padding(pe_long)

pe_long <- pe_long %>%
  dplyr::mutate(
    minute_start = (pe_window - 1L) * bin_minutes,
    minute_mid = minute_start + bin_minutes / 2,
    hour_mid = minute_mid / 60,
    day_mid = hour_mid / 24)

pe_long <- pe_long %>%
  dplyr::rename(FlyID = FlyID_key)

#### Truncate to 14 days to align with activity pipeline, remove _Control flies from second block ####

pe_long_14d <- pe_long %>%
  filter(pe_window <= max_windows_14d)

# confirm
pe_long_14d %>%
  summarise(max_window = max(pe_window), max_minute_start = max(minute_start)) %>%
  print()
```

```
## # A tibble: 1 × 2
##   max_window max_minute_start
##        <int>            <int>
## 1        672            20130
```

```
pe_long_14d <- pe_long_14d %>% filter(!str_detect(FlyID, "_Control"))

#### Join true class labels (alive/dead) by FlyID ####

# Use the SAME analysed fly data as the activity pipeline
# Build reference class table from the analysis dataset
# use infected_df because it already excludes controls (UI) 
stopifnot(exists("infected_df") || exists("fly_data_pre"))

ref_class_clean <- if (exists("infected_df")) {
  infected_df %>%
    distinct(FlyID, class) %>%
    mutate(
      FlyID = as.character(FlyID),
      class = tolower(as.character(class))
    )
} else {
  # build from fly_data_pre then filter to infected
  fly_data_pre %>%
    filter(Treatment != "UI") %>%
    distinct(FlyID, class) %>%
    mutate(
      FlyID = as.character(FlyID),
      class = tolower(as.character(class))
    )
}

# should be one class per fly
ref_class_clean <- ref_class_clean %>%
  group_by(FlyID) %>%
  summarise(class = first(class), .groups = "drop")

valid_ids <- ref_class_clean$FlyID

# check which PE IDs won't be used for diagnostics - this should only show flies dropped because they were controls or because they died before 600 minute mark
pe_only_ids <- setdiff(unique(pe_long_14d$FlyID), valid_ids)
if (length(pe_only_ids) > 0) {
  message("Dropping PE-only flies not present in analysed activity set (showing up to 200): ",
          paste(head(pe_only_ids, 200), collapse = ", "))
}
```

```
## Dropping PE-only flies not present in analysed activity set (showing up to 200): M14T6_G3Dead, M15T22_C4Dead, M16T25_P4Dead, M16T31_P3Dead, M18T30_P4Dead, M19T8_C4Dead, M19T28_G1Dead, DeadM12T22, DeadM13T5, DeadM13T27, DeadM14T3, DeadM14T4, DeadM14T12, DeadM14T19, DeadM14T27, DeadM15T7, DeadM15T18, DeadM15T27, DeadM16T7, DeadM16T12, DeadM16T24, DeadM16T25, DeadM17T15, DeadM17T16, DeadM18T31, DeadM19T1, DeadM19T25
```

```
# Restrict and join 
pe_long_14d <- pe_long_14d %>%
  filter(FlyID %in% valid_ids) %>%
  left_join(ref_class_clean, by = "FlyID")

# fail if anything still didn't join
if (any(is.na(pe_long_14d$class))) {
  missing_ids <- pe_long_14d %>% filter(is.na(class)) %>% distinct(FlyID)
  stop("PE FlyIDs still missing class after join. IDs: ",
       paste(head(missing_ids$FlyID, 25), collapse = ", "))
}

#### ANALYSES and FIGURES ####

##### Mean PE per fly: over full 14-day period after trimming #####
# The unit of replication is fly not time point!!!!!!

fly_total <- pe_long_14d %>%
  mutate(block = case_when(
    stringr::str_detect(source_file, "Monitor") ~ "blockA_monitor",
    stringr::str_detect(source_file, "Kabir")   ~ "blockB_kabir", 
    TRUE ~ NA_character_)) %>%
  filter(!is.na(block)) %>%
  group_by(FlyID, class, block) %>%
  summarise(total_mean_entropy = mean(entropy, na.rm = TRUE), .groups = "drop") %>%
  mutate(class = factor(class, levels = c("alive", "dead")), 
         block = factor(block))

head(fly_total)
```

```
## # A tibble: 6 × 4
##   FlyID       class block        total_mean_entropy
##   <chr>       <fct> <fct>                     <dbl>
## 1 AliveM12T11 alive blockB_kabir              1.30 
## 2 AliveM12T19 alive blockB_kabir              0.790
## 3 AliveM12T26 alive blockB_kabir              1.28 
## 4 AliveM12T27 alive blockB_kabir              1.57 
## 5 AliveM12T28 alive blockB_kabir              1.45 
## 6 AliveM12T5  alive blockB_kabir              1.38
```

```
str(fly_total)
```

```
## tibble [411 × 4] (S3: tbl_df/tbl/data.frame)
##  $ FlyID             : chr [1:411] "AliveM12T11" "AliveM12T19" "AliveM12T26" "AliveM12T27" ...
##  $ class             : Factor w/ 2 levels "alive","dead": 1 1 1 1 1 1 1 1 1 1 ...
##  $ block             : Factor w/ 2 levels "blockA_monitor",..: 2 2 2 2 2 2 2 2 2 2 ...
##  $ total_mean_entropy: num [1:411] 1.3 0.79 1.28 1.57 1.45 ...
```

```
##### What are the class means? #####
class_means <- fly_total %>%
  group_by(class) %>%
  summarise(
    mean_entropy = mean(total_mean_entropy, na.rm = TRUE),
    sd_entropy   = sd(total_mean_entropy, na.rm = TRUE),
    n            = n(),
    .groups = "drop")

class_means
```

```
## # A tibble: 2 × 4
##   class mean_entropy sd_entropy     n
##   <fct>        <dbl>      <dbl> <int>
## 1 alive         1.27      0.249   161
## 2 dead          1.21      0.624   250
```

```
##### Plot mean entropy #####
p_mean_ent <- ggplot(fly_total, aes(x = class, y = total_mean_entropy, fill = class)) +
  geom_boxplot(outlier.shape = NA, alpha = 0.8, na.rm = TRUE, position = position_dodge()) +
  geom_point(na.rm = TRUE, position = position_jitterdodge(0.07), pch=21) +
  scale_y_continuous(limits = c(0, 2.7), breaks = c(0, 0.5, 1, 1.5, 2, 2.5)) +
  scale_fill_manual(values = c("#0072B2","#D55E00")) +
  labs(x = "Class", y = "Mean PE (total)") +
  theme_light() +
  theme(legend.position = "none",
        axis.text = element_text(size=14),
        axis.title = element_text(size = 16)); p_mean_ent
```

```
##### Mixed model: test difference in PE mean total between dead and alive flies #####
###### lmer model (Assumes equal variance) ######
m1 <- lmer(total_mean_entropy ~ class + (1|block), data = fly_total, REML = TRUE)
summary(m1)
```

```
## Linear mixed model fit by REML. t-tests use Satterthwaite's method [
## lmerModLmerTest]
## Formula: total_mean_entropy ~ class + (1 | block)
##    Data: fly_total
## 
## REML criterion at convergence: 620.5
## 
## Scaled residuals: 
##      Min       1Q   Median       3Q      Max 
## -2.29612 -0.52492  0.09213  0.63990  2.19624 
## 
## Random effects:
##  Groups   Name        Variance Std.Dev.
##  block    (Intercept) 0.004312 0.06567 
##  Residual             0.259144 0.50906 
## Number of obs: 411, groups:  block, 2
## 
## Fixed effects:
##              Estimate Std. Error        df t value Pr(>|t|)   
## (Intercept)   1.26869    0.06138   1.82534  20.668  0.00355 **
## classdead    -0.06192    0.05147 408.24045  -1.203  0.22964   
## ---
## Signif. codes:  0 '***' 0.001 '**' 0.01 '*' 0.05 '.' 0.1 ' ' 1
## 
## Correlation of Fixed Effects:
##           (Intr)
## classdead -0.510
```

```
car::Anova(m1, type = 2)
```

```
## Analysis of Deviance Table (Type II Wald chisquare tests)
## 
## Response: total_mean_entropy
##        Chisq Df Pr(>Chisq)
## class 1.4474  1     0.2289
```

```
check_model(m1)
```

```
plot(m1)
```

```
###### lme model accounting for different variance between classes (varIdent) ######
m1_var <- lme(total_mean_entropy ~ class, random  = ~ 1|block, 
              weights = varIdent(form = ~ 1|class), data = fly_total, method = "REML")
summary(m1_var)
```

```
## Linear mixed-effects model fit by REML
##   Data: fly_total 
##        AIC     BIC    logLik
##   501.3404 521.409 -245.6702
## 
## Random effects:
##  Formula: ~1 | block
##         (Intercept) Residual
## StdDev:  0.03346828 0.245399
## 
## Variance function:
##  Structure: Different standard deviations per stratum
##  Formula: ~1 | class 
##  Parameter estimates:
##    alive     dead 
## 1.000000 2.560308 
## Fixed effects:  total_mean_entropy ~ class 
##                  Value  Std.Error  DF  t-value p-value
## (Intercept)  1.2727801 0.03057815 408 41.62384  0.0000
## classdead   -0.0664929 0.04420654 408 -1.50414  0.1333
##  Correlation: 
##           (Intr)
## classdead -0.277
## 
## Standardized Within-Group Residuals:
##        Min         Q1        Med         Q3        Max 
## -4.2156360 -0.6051211  0.1057552  0.7387933  2.1814260 
## 
## Number of Observations: 411
## Number of Groups: 2
```

```
plot(m1_var)
```

```
check_model(m1_var)
```

```
## Converting missing values (`NA`) into regular values currently not
##   possible for variables of class `NULL`.
```

```
qqnorm(resid(m1_var)); qqline(resid(m1_var))
```

```
car::Anova(m1_var, type = c(2))
```

```
## Analysis of Deviance Table (Type II tests)
## 
## Response: total_mean_entropy
##        Chisq Df Pr(>Chisq)
## class 2.2624  1     0.1325
```

```
##### Mean PE per fly over 0–20 hpi #####
# ACUTE INFECTION
# (0–20 hours = 1200 minutes)

fly_total_0_20 <- pe_long_14d %>%
  mutate(block = case_when( 
    stringr::str_detect(source_file, "Monitor") ~ "blockA_monitor", 
    stringr::str_detect(source_file, "Kabir")   ~ "blockB_kabir",
      TRUE ~ NA_character_)) %>%
  filter(!is.na(block)) %>%
  filter(minute_start >= 0, minute_start < 20 * 60) %>%
  group_by(FlyID, block, class) %>%
  summarise(
    total_mean_entropy = mean(entropy, na.rm = TRUE),
    .groups = "drop") %>%
  mutate(
    class = factor(class, levels = c("alive", "dead")),
    block = factor(block))

##### non parametric tests: not needed, but will leave if useful #####
##### Without block #####
print(wilcox.test(total_mean_entropy ~ class, data = fly_total_0_20))
```

```
## 
##  Wilcoxon rank sum test with continuity correction
## 
## data:  total_mean_entropy by class
## W = 24755, p-value = 8.211e-05
## alternative hypothesis: true location shift is not equal to 0
```

```
##### With block #####
p_blocked_pe020 <- wilcox_test(total_mean_entropy ~ class|block, data = fly_total_0_20, distribution = "approximate")
p_blocked_pe020
```

```
## 
##  Approximative Wilcoxon-Mann-Whitney Test
## 
## data:  total_mean_entropy by
##   class (alive, dead) 
##   stratified by block
## Z = 3.8715, p-value < 1e-04
## alternative hypothesis: true mu is not equal to 0
```

```
# Cliff's delta effect size
if (requireNamespace("effectsize", quietly = TRUE)) {
  cd_020 <- effectsize::cliffs_delta(total_mean_entropy ~ class, data = fly_total_0_20)
  print(cd_020)}
```

```
## r (rank biserial) |       95% CI
## --------------------------------
## 0.23              | [0.12, 0.34]
```

```
# Bootstrap CI for median difference (alive - dead)
set.seed(1)
B <- 2000
alive_vals <- fly_total_0_20$total_mean_entropy[fly_total_0_20$class == "alive"]
dead_vals  <- fly_total_0_20$total_mean_entropy[fly_total_0_20$class == "dead"]

boot_med_diff <- replicate(B,
                           median(sample(alive_vals, replace = TRUE)) - median(sample(dead_vals, replace = TRUE)))
print(quantile(boot_med_diff, c(0.025, 0.5, 0.975), na.rm = TRUE))
```

```
##      2.5%       50%     97.5% 
## 0.1171918 0.2156058 0.3429343
```

```
##### Boxplot - differences ACUTE PHASE (0-20: when most flies die) #####
p_mean_ent_020 <- ggplot(fly_total_0_20, aes(x = class, y = total_mean_entropy, fill = class)) +
  geom_boxplot(outlier.shape = NA, alpha = 0.8, na.rm = TRUE, position = position_dodge()) +
  geom_point(na.rm = TRUE, position = position_jitterdodge(0.07), pch=21) +
  scale_fill_manual(values = c("#0072B2","#D55E00")) +
  labs(x = "Class", y = "Mean permutation entropy (0–20 hpi)") +
  theme_light() +
  theme(legend.position = "none",
        axis.text = element_text(size=14),
        axis.title = element_text(size = 16)); p_mean_ent_020
```

```
###### Mixed model PE 0-20 hpi ######
m2 <- lmer(total_mean_entropy ~ class + (1|block), data = fly_total_0_20, REML = TRUE)
summary(m2)
```

```
## Linear mixed model fit by REML. t-tests use Satterthwaite's method [
## lmerModLmerTest]
## Formula: total_mean_entropy ~ class + (1 | block)
##    Data: fly_total_0_20
## 
## REML criterion at convergence: 503.2
## 
## Scaled residuals: 
##     Min      1Q  Median      3Q     Max 
## -3.3438 -0.6649  0.1536  0.7172  1.9056 
## 
## Random effects:
##  Groups   Name        Variance Std.Dev.
##  block    (Intercept) 0.001678 0.04096 
##  Residual             0.194772 0.44133 
## Number of obs: 411, groups:  block, 2
## 
## Fixed effects:
##              Estimate Std. Error        df t value Pr(>|t|)    
## (Intercept)   1.72801    0.04528   2.43043  38.164 0.000195 ***
## classdead    -0.18132    0.04462 408.36799  -4.064 5.79e-05 ***
## ---
## Signif. codes:  0 '***' 0.001 '**' 0.01 '*' 0.05 '.' 0.1 ' ' 1
## 
## Correlation of Fixed Effects:
##           (Intr)
## classdead -0.600
```

```
car::Anova(m2, type = 2)
```

```
## Analysis of Deviance Table (Type II Wald chisquare tests)
## 
## Response: total_mean_entropy
##        Chisq Df Pr(>Chisq)    
## class 16.515  1  4.827e-05 ***
## ---
## Signif. codes:  0 '***' 0.001 '**' 0.01 '*' 0.05 '.' 0.1 ' ' 1
```

```
check_model(m2)
```

```
plot(m2)
```

```
##### PE time series, x-axis in HOURS #####
# Can also be analysed using Generalised Additive Models - did prelim analysis in October 

pe_mean_ts <- pe_long_14d %>%
  group_by(class, hour_mid) %>%
  summarise(mean_entropy = mean(entropy, na.rm = TRUE), .groups = "drop")

p_pe_ts <- ggplot() +
  geom_line(data = pe_long_14d, aes(x = hour_mid, y = entropy, group = FlyID, color = class), alpha = 0.08) +
  geom_line(data = pe_mean_ts, aes(x = hour_mid, y = mean_entropy), linewidth = 1.0, color = "black") +
  facet_wrap(~class, ncol = 1) +
  labs(x = "Hours post infection (30 min windows)",
       y = "Permutation entropy (PE)") +
  scale_y_continuous(limits = c(0, 2.7), breaks = c(0, 0.5, 1, 1.5, 2, 2.5, 3)) +
  scale_x_continuous(limits = c(0, 337), breaks = c(0, 25, 50, 75, 100, 125, 150, 175, 200, 225, 250, 275, 300, 325)) +
  coord_cartesian(expand = c(0,0)) +
  theme_minimal() +
  theme(legend.position = "none",
        axis.text = element_text(size=14),
        axis.title = element_text(size = 16),
        strip.text.x = element_text(size = 14)) +
  theme(legend.position = "none") +
  scale_color_manual(values = c("alive" = "#0072B2", "dead" = "#D55E00")); p_pe_ts
```

```
##### Early vs late acute phase windows (no pseudoreplication) #####
# Define windows: 0–4 hpi and 16–20 hpi
# - per-fly mean in each window
# - paired Wilcoxon within class (alive early vs alive late; dead early vs dead late)
# - unpaired Wilcoxon between classes within a window (alive vs dead at early; alive vs dead at late)
# - mixed model controlling for two experimental blocks - USE THIS ANALYSIS!

pe_win <- pe_long_14d %>%
  mutate(
    window = case_when(
      minute_start >= 0       & minute_start <  4*60  ~ "early",
      minute_start >= 12*60   & minute_start < 16*60  ~ "late", # minute_start >= 16*60   & minute_start < 20*60  ~ "late"
      TRUE ~ NA_character_),
    block = case_when(
      str_detect(source_file, "Monitor") ~ "blockA_monitor",
      str_detect(source_file, "Kabir") ~ "blockB_kabir",
      TRUE ~ NA_character_)) %>%
  filter(!is.na(window), !is.na(block))

fly_means <- pe_win %>%
  group_by(FlyID, block, class, window) %>%
  summarise(mean_entropy = mean(entropy, na.rm = TRUE), .groups = "drop") %>%
  mutate(
    window = factor(window, levels = c("early", "late")),
    class  = factor(class,  levels = c("alive", "dead")),
    block  = factor(block))

# counts contributing per window
print(fly_means %>% distinct(FlyID, class, window) %>% count(class, window))
```

```
## # A tibble: 4 × 3
##   class window     n
##   <fct> <fct>  <int>
## 1 alive early    161
## 2 alive late     161
## 3 dead  early    250
## 4 dead  late     249
```

```
# Paired early vs late within alive
alive_wide <- fly_means %>%
  filter(class == "alive") %>%
  select(FlyID, window, mean_entropy) %>%
  pivot_wider(names_from = window, values_from = mean_entropy) %>%
  filter(!is.na(early), !is.na(late))

alive_test <- wilcox.test(alive_wide$early, alive_wide$late, paired = TRUE)
print(alive_test)
```

```
## 
##  Wilcoxon signed rank test with continuity correction
## 
## data:  alive_wide$early and alive_wide$late
## V = 9108, p-value = 1.263e-05
## alternative hypothesis: true location shift is not equal to 0
```

```
# Paired early vs late within dead
dead_wide <- fly_means %>%
  filter(class == "dead") %>%
  select(FlyID, window, mean_entropy) %>%
  pivot_wider(names_from = window, values_from = mean_entropy) %>%
  filter(!is.na(early), !is.na(late))

dead_test <- wilcox.test(dead_wide$early, dead_wide$late, paired = TRUE)
print(dead_test)
```

```
## 
##  Wilcoxon signed rank test with continuity correction
## 
## data:  dead_wide$early and dead_wide$late
## V = 22368, p-value = 2.21e-09
## alternative hypothesis: true location shift is not equal to 0
```

```
# Optional paired effect sizes in rstatix
if (requireNamespace("rstatix", quietly = TRUE)) {
  alive_eff <- alive_wide %>%
    mutate(id = row_number()) %>%
    pivot_longer(cols = c(early, late), names_to = "phase", values_to = "mean_entropy") %>%
    mutate(phase = factor(phase, levels = c("early", "late"))) %>%
    rstatix::wilcox_effsize(mean_entropy ~ phase, paired = TRUE)
  print(alive_eff)
  
  dead_eff <- dead_wide %>%
    mutate(id = row_number()) %>%
    pivot_longer(cols = c(early, late), names_to = "phase", values_to = "mean_entropy") %>%
    mutate(phase = factor(phase, levels = c("early", "late"))) %>%
    rstatix::wilcox_effsize(mean_entropy ~ phase, paired = TRUE)
  print(dead_eff)}
```

```
## # A tibble: 1 × 7
##   .y.          group1 group2 effsize    n1    n2 magnitude
## * <chr>        <chr>  <chr>    <dbl> <int> <int> <ord>    
## 1 mean_entropy early  late     0.344   161   161 moderate 
## # A tibble: 1 × 7
##   .y.          group1 group2 effsize    n1    n2 magnitude
## * <chr>        <chr>  <chr>    <dbl> <int> <int> <ord>    
## 1 mean_entropy early  late     0.379   249   249 moderate
```

```
# Median paired differences (late - early)
alive_delta <- median(alive_wide$late - alive_wide$early, na.rm = TRUE)
dead_delta  <- median(dead_wide$late  - dead_wide$early,  na.rm = TRUE)
print(c(alive_delta = alive_delta, dead_delta = dead_delta))
```

```
## alive_delta  dead_delta 
##  -0.1394964  -0.1700950
```

```
# Between-class comparisons within each window
early_df <- fly_means %>% filter(window == "early")
late_df  <- fly_means %>% filter(window == "late")

print(wilcox.test(mean_entropy ~ class, data = early_df))
```

```
## 
##  Wilcoxon rank sum test with continuity correction
## 
## data:  mean_entropy by class
## W = 23991, p-value = 0.001008
## alternative hypothesis: true location shift is not equal to 0
```

```
print(wilcox.test(mean_entropy ~ class, data = late_df))
```

```
## 
##  Wilcoxon rank sum test with continuity correction
## 
## data:  mean_entropy by class
## W = 24517, p-value = 0.0001354
## alternative hypothesis: true location shift is not equal to 0
```

```
##### Blocked comparison #####
p_blocked_early <- wilcox_test(mean_entropy ~ class|block, data = early_df, distribution = "approximate")
p_blocked_early
```

```
## 
##  Approximative Wilcoxon-Mann-Whitney Test
## 
## data:  mean_entropy by
##   class (alive, dead) 
##   stratified by block
## Z = 3.4195, p-value = 7e-04
## alternative hypothesis: true mu is not equal to 0
```

```
p_blocked_late <- wilcox_test(mean_entropy ~ class|block, data = late_df, distribution = "approximate")
p_blocked_late
```

```
## 
##  Approximative Wilcoxon-Mann-Whitney Test
## 
## data:  mean_entropy by
##   class (alive, dead) 
##   stratified by block
## Z = 3.7602, p-value = 4e-04
## alternative hypothesis: true mu is not equal to 0
```

```
##### Effect sizes #####
# Cliff's delta by window (optional)
if (requireNamespace("effectsize", quietly = TRUE)) {
  print(effectsize::cliffs_delta(mean_entropy ~ class, data = early_df))
  print(effectsize::cliffs_delta(mean_entropy ~ class, data = late_df))
}
```

```
## r (rank biserial) |       95% CI
## --------------------------------
## 0.19              | [0.08, 0.30]
## r (rank biserial) |       95% CI
## --------------------------------
## 0.22              | [0.11, 0.33]
```

```
##### Plots for early/late window comparisons #####

# "alive early", "alive late", "dead early", "dead late"
fly_means_named <- fly_means %>%
  mutate(
    new_class = paste0(as.character(class), " ", as.character(window)),
    new_class = factor(new_class, levels = c("alive early","alive late","dead early","dead late")))

##### Early vs late acute phase Box plot ######
# early vs late on x; class as color/fill + means connected
plot_df <- fly_means %>%
  mutate(window = factor(window, levels = c("early","late")),
         class  = factor(class, levels = c("alive","dead")))

mean_df <- plot_df %>%
  group_by(class, window) %>%
  summarise(mean_entropy = mean(mean_entropy, na.rm = TRUE), .groups = "drop")

pd <- position_dodge(width = 0.55)

p_mean_ent_020 <- ggplot(fly_total_0_20, aes(x = class, y = total_mean_entropy, fill = class)) +
  geom_boxplot(outlier.shape = NA, alpha = 0.8, na.rm = TRUE, position = position_dodge()) +
  geom_point(na.rm = TRUE, position = position_jitterdodge(0.07), pch=21) +
  scale_fill_manual(values = c("#0072B2","#D55E00")) +
  labs(x = "Class", y = "Permutation entropy (0–20 hpi)") +
  theme_light() +
  theme(legend.position = "none",
        axis.text = element_text(size=14),
        axis.title = element_text(size = 16)); p_mean_ent_020
```

```
# Consistent ordering and colours, classes were confused in the original rf confusion matrix and no one saw this!!
plot_df$class <- factor(plot_df$class, levels = c("alive","dead"))
mean_df$class <- factor(mean_df$class, levels = c("alive","dead"))

pal_class <- c(alive = "#0072B2", dead = "#D55E00")

pd <- position_dodge(width = 0.8)

p_fly2 <- ggplot(plot_df, aes(x = window, y = mean_entropy)) +
  geom_boxplot(aes(fill = class), alpha = 0.8, na.rm = TRUE, position = pd, outlier.shape = NA) +
  geom_point(
    aes(fill = class),                  
    na.rm = TRUE,
    position = position_jitterdodge(jitter.width = 0.07, dodge.width = 0.8),
    pch = 21,
    size = 1.7,
    colour = "black",
    stroke = 0.2) +
  # mean lines
  geom_line(
    data = mean_df,
    aes(y = mean_entropy, group = class),
    position = pd, linewidth = 1.2, alpha = 0.7, 
    colour = "black") +
  # mean points 
  geom_point(
    data = mean_df,
    aes(y = mean_entropy, group = class), 
    position = pd,
    size = 3,
    colour = "black") + 
  scale_fill_manual(values = pal_class, breaks = c("alive","dead")) +
  scale_colour_manual(values = pal_class, breaks = c("alive","dead")) +
  scale_x_discrete(labels = c("early" = "0-4", "late" = "12-16")) +
  labs(x = "Hours post infection",
       y = "Mean PE", 
       colour = "Class", 
       fill = "Class") +
  theme_light(base_size = 12) +
  theme(panel.grid.minor = element_blank(), 
        axis.text = element_text(size = 14),
        axis.title = element_text(size = 16),
        legend.title = element_text(size = 14),
        legend.text  = element_text(size = 12), 
        legend.position = "none"); p_fly2
```

```
## Ignoring unknown labels:
## • colour : "Class"
```

```
## Warning: No shared levels found between `names(values)` of the manual scale and the
## data's colour values.
```

```
##### Mixed model for early/late acute period (single analysis controlling for FlyID and block) #####

# NB non-parametric tests are not needed if using mixed models

# NB the fit is singular only because block is used as a random effect. Individual flyIDs contribute all of the variance.

if (requireNamespace("lme4", quietly = TRUE) && requireNamespace("lmerTest", quietly = TRUE) && 
    requireNamespace("performance", quietly = TRUE)) {
  m3 <- lmer(mean_entropy ~ class * window + (1|block/FlyID), data = fly_means, REML = TRUE)
  print(summary(m3))
  print(car::Anova(m3, type = 2))
  print(check_model(m3))
  plot(m3)
  qqnorm(resid(m3)); qqline(resid(m3))}
```

```
## Linear mixed model fit by REML. t-tests use Satterthwaite's method [
## lmerModLmerTest]
## Formula: mean_entropy ~ class * window + (1 | block/FlyID)
##    Data: fly_means
## 
## REML criterion at convergence: 1215.3
## 
## Scaled residuals: 
##     Min      1Q  Median      3Q     Max 
## -2.9433 -0.4304  0.1500  0.5018  2.2651 
## 
## Random effects:
##  Groups      Name        Variance  Std.Dev.
##  FlyID:block (Intercept) 0.1577834 0.39722 
##  block       (Intercept) 0.0005543 0.02354 
##  Residual                0.1398259 0.37393 
## Number of obs: 821, groups:  FlyID:block, 411; block, 2
## 
## Fixed effects:
##                       Estimate Std. Error        df t value Pr(>|t|)    
## (Intercept)            2.07167    0.04611   6.48098  44.925 2.49e-09 ***
## classdead             -0.16766    0.05514 636.94341  -3.041  0.00246 ** 
## windowlate            -0.11929    0.04168 407.45550  -2.862  0.00442 ** 
## classdead:windowlate  -0.08568    0.05347 407.68604  -1.602  0.10985    
## ---
## Signif. codes:  0 '***' 0.001 '**' 0.01 '*' 0.05 '.' 0.1 ' ' 1
## 
## Correlation of Fixed Effects:
##             (Intr) clssdd wndwlt
## classdead   -0.727              
## windowlate  -0.452  0.378       
## clssdd:wndw  0.352 -0.484 -0.779
## Analysis of Deviance Table (Type II Wald chisquare tests)
## 
## Response: mean_entropy
##                Chisq Df Pr(>Chisq)    
## class        19.0326  1  1.285e-05 ***
## window       43.0654  1  5.294e-11 ***
## class:window  2.5676  1     0.1091    
## ---
## Signif. codes:  0 '***' 0.001 '**' 0.01 '*' 0.05 '.' 0.1 ' ' 1
```

```
sim_m3<- simulateResiduals(m3, n=1000)
plot(sim_m3)
testDispersion(sim_m3)
```

```
## 
##  DHARMa nonparametric dispersion test via sd of residuals fitted vs.
##  simulated
## 
## data:  simulationOutput
## dispersion = 0.99604, p-value = 0.936
## alternative hypothesis: two.sided
```

```
testUniformity(sim_m3)
```

```
## 
##  Asymptotic one-sample Kolmogorov-Smirnov test
## 
## data:  simulationOutput$scaledResiduals
## D = 0.12479, p-value = 1.571e-11
## alternative hypothesis: two-sided
```

```
##### Model unequal variances with lme, see Zuur et al mixed model book for more detail, also Kutzer et al 2016 Ecology and Evolution #####
# Model accounting for unequal variances in class
m3_var <- lme(mean_entropy ~ class * window, random  = ~ 1|block/FlyID, 
              weights = varIdent(form = ~ 1|class), data = fly_means, method = "REML")
summary(m3_var)
```

```
## Linear mixed-effects model fit by REML
##   Data: fly_means 
##        AIC      BIC    logLik
##   1220.556 1258.201 -602.2779
## 
## Random effects:
##  Formula: ~1 | block
##         (Intercept)
## StdDev:  0.02323583
## 
##  Formula: ~1 | FlyID %in% block
##         (Intercept)  Residual
## StdDev:   0.3904904 0.4076838
## 
## Variance function:
##  Structure: Different standard deviations per stratum
##  Formula: ~1 | class 
##  Parameter estimates:
##      dead     alive 
## 1.0000000 0.7958051 
## Fixed effects:  mean_entropy ~ class * window 
##                           Value  Std.Error  DF  t-value p-value
## (Intercept)           2.0716576 0.04326287 408 47.88535  0.0000
## classdead            -0.1676518 0.05363477 408 -3.12580  0.0019
## windowlate           -0.1192862 0.03616032 408 -3.29881  0.0011
## classdead:windowlate -0.0854939 0.05139227 408 -1.66355  0.0970
##  Correlation: 
##                      (Intr) clssdd wndwlt
## classdead            -0.690              
## windowlate           -0.418  0.337       
## classdead:windowlate  0.294 -0.478 -0.704
## 
## Standardized Within-Group Residuals:
##        Min         Q1        Med         Q3        Max 
## -2.9706268 -0.4432360  0.1463530  0.5175974  2.0358242 
## 
## Number of Observations: 821
## Number of Groups: 
##            block FlyID %in% block 
##                2              411
```

```
plot(m3_var)
```

```
qqnorm(resid(m3_var)); qqline(resid(m3_var))
```

```
car::Anova(m3_var, type = c(2))     # fixed effects tests
```

```
## Analysis of Deviance Table (Type II tests)
## 
## Response: mean_entropy
##                Chisq Df Pr(>Chisq)    
## class        19.9425  1  7.981e-06 ***
## window       39.5597  1  3.182e-10 ***
## class:window  2.7674  1     0.0962 .  
## ---
## Signif. codes:  0 '***' 0.001 '**' 0.01 '*' 0.05 '.' 0.1 ' ' 1
```

```
# Create an interaction column to model variance within class:window
fly_means <- fly_means %>%
  dplyr::mutate(class_window = interaction(class, window, drop = TRUE))

table(fly_means$class_window, useNA = "ifany")
```

```
## 
## alive.early  dead.early  alive.late   dead.late 
##         161         250         161         249
```

```
# Fit model with class:window
m3_var_cw <- lme(mean_entropy ~ class * window, random = ~ 1 | block/FlyID, 
                 weights = varIdent(form = ~ 1 | class_window), data = fly_means, method = "REML")

summary(m3_var_cw)
```

```
## Linear mixed-effects model fit by REML
##   Data: fly_means 
##        AIC      BIC   logLik
##   1215.136 1262.192 -597.568
## 
## Random effects:
##  Formula: ~1 | block
##         (Intercept)
## StdDev:  0.01506334
## 
##  Formula: ~1 | FlyID %in% block
##         (Intercept)  Residual
## StdDev:   0.3906312 0.3650379
## 
## Variance function:
##  Structure: Different standard deviations per stratum
##  Formula: ~1 | class_window 
##  Parameter estimates:
##  dead.early   dead.late alive.early  alive.late 
##   1.0000000   1.2212713   1.0520343   0.6938187 
## Fixed effects:  mean_entropy ~ class * window 
##                           Value  Std.Error  DF  t-value p-value
## (Intercept)           2.0713164 0.04447134 408 46.57644  0.0000
## classdead            -0.1672706 0.05484284 408 -3.05000  0.0024
## windowlate           -0.1192862 0.03625536 408 -3.29017  0.0011
## classdead:windowlate -0.0856819 0.05144691 408 -1.66544  0.0966
##  Correlation: 
##                      (Intr) clssdd wndwlt
## classdead            -0.764              
## windowlate           -0.568  0.461       
## classdead:windowlate  0.400 -0.514 -0.705
## 
## Standardized Within-Group Residuals:
##        Min         Q1        Med         Q3        Max 
## -3.0101634 -0.4418920  0.1394151  0.5335372  2.1528701 
## 
## Number of Observations: 821
## Number of Groups: 
##            block FlyID %in% block 
##                2              411
```

```
car::Anova(m3_var_cw, type = 2)
```

```
## Analysis of Deviance Table (Type II tests)
## 
## Response: mean_entropy
##                Chisq Df Pr(>Chisq)    
## class        20.7151  1  5.329e-06 ***
## window       39.5841  1  3.142e-10 ***
## class:window  2.7737  1    0.09582 .  
## ---
## Signif. codes:  0 '***' 0.001 '**' 0.01 '*' 0.05 '.' 0.1 ' ' 1
```

```
plot(m3_var_cw)
```

```
qqnorm(resid(m3_var_cw)); qqline(resid(m3_var_cw))
```

```
# Fit model accounting for variance in window only
m3_base <- lme(mean_entropy ~ class * window, random = ~1|block/FlyID, 
               data = fly_means, method="REML")
m3_var_window <- update(m3_base, weights = varIdent(form = ~1|window))
summary(m3_var_window)
```

```
## Linear mixed-effects model fit by REML
##   Data: fly_means 
##       AIC      BIC    logLik
##   1230.94 1268.585 -607.4699
## 
## Random effects:
##  Formula: ~1 | block
##         (Intercept)
## StdDev:  0.03023763
## 
##  Formula: ~1 | FlyID %in% block
##         (Intercept)  Residual
## StdDev:   0.3971411 0.3630405
## 
## Variance function:
##  Structure: Different standard deviations per stratum
##  Formula: ~1 | window 
##  Parameter estimates:
##    early     late 
## 1.000000 1.059097 
## Fixed effects:  mean_entropy ~ class * window 
##                           Value  Std.Error  DF  t-value p-value
## (Intercept)           2.0719927 0.04750352 408 43.61767  0.0000
## classdead            -0.1680262 0.05438561 408 -3.08953  0.0021
## windowlate           -0.1192862 0.04167568 408 -2.86225  0.0044
## classdead:windowlate -0.0857394 0.05346897 408 -1.60354  0.1096
##  Correlation: 
##                      (Intr) clssdd wndwlt
## classdead            -0.697              
## windowlate           -0.414  0.361       
## classdead:windowlate  0.322 -0.463 -0.779
## 
## Standardized Within-Group Residuals:
##        Min         Q1        Med         Q3        Max 
## -2.9506265 -0.4338401  0.1403661  0.5036468  2.2992091 
## 
## Number of Observations: 821
## Number of Groups: 
##            block FlyID %in% block 
##                2              411
```

```
# AIC of each model, which has lowest?
AIC(m3_base, m3_var, m3_var_window, m3_var_cw)
```

```
##               df      AIC
## m3_base        7 1229.327
## m3_var         8 1220.556
## m3_var_window  8 1230.940
## m3_var_cw     10 1215.136
```

```
# Importantly, results don't change, but will keep correct variance structure!

###### Adjust p-values in m1, m2 and m3 for multiple testing ######
# Main effects
a1 <- car::Anova(m1, type = 2)  # class
a2 <- car::Anova(m2, type = 2)  # class
a3 <- car::Anova(m3_var_cw, type = 2)  # class, window, class:window

# For reuse in future!, extract p-values for main effects and/or interactions
p_raw <- c(
  m1_class = a1["class", "Pr(>Chisq)"] %||% a1["class", "Pr(>F)"],
  m2_class = a2["class", "Pr(>Chisq)"] %||% a2["class", "Pr(>F)"],
  m3_class = a3["class", "Pr(>Chisq)"] %||% a3["class", "Pr(>F)"],
  m3_window = a3["window", "Pr(>Chisq)"] %||% a3["window", "Pr(>F)"],
  m3_class_window = a3["class:window", "Pr(>Chisq)"] %||% a3["class:window", "Pr(>F)"])

# View the raw p-values (no correction)
p_raw
```

```
##        m1_class        m2_class        m3_class       m3_window m3_class_window 
##    2.289396e-01    4.826607e-05    5.329298e-06    3.142366e-10    9.582436e-02
```

```
p_adj <- p.adjust(p_raw, method = "BH") # change method if needed: BH = Benjamini-Hochberg; bonferroni = Bonferroni correction

# View the adjusted p-values
p_adj
```

```
##        m1_class        m2_class        m3_class       m3_window m3_class_window 
##    2.289396e-01    8.044345e-05    1.332325e-05    1.571183e-09    1.197805e-01
```

```
#### Permutation entropy figure for ms ####
right_column <- plot_grid(
  p_mean_ent, p_fly2,
  ncol = 1,
  labels = c("B", "C"),
  label_fontface = "bold",
  label_size = 16,
  label_x = 0.02, 
  label_y = 0.98)
```

```
## Ignoring unknown labels:
## • colour : "Class"
```

```
## Warning: No shared levels found between `names(values)` of the manual scale and the
## data's colour values.
```

```
final_pe_figure <- plot_grid(
  p_pe_ts, right_column,
  ncol = 2,
  rel_widths = c(2/3, 1/3),
  labels = c("A", ""), # label only the left block
  label_fontface = "bold",
  label_size = 16,
  label_x = 0.02,
  label_y = 0.98)

final_pe_figure
```

```
# Save figure 
save_plot("/Users/mkutzer/Desktop/Critical transitions/Manuscript/Figure_3.png", base_width = 10, base_height = 6, final_pe_figure)
```
